# Spinal Noradrenergic Overdrive Promotes Reversible Motor Unit Aging and Sarcopenia

**DOI:** 10.64898/2026.10.30.763089

**Authors:** Zhong-Min Wang, Valentina Grinevich, María Laura Messi, Savannah Longo, Ragunath Muthu, William Ryan Meeker, Evgeny Budygin, Osvaldo Delbono

## Abstract

**Background:** Sarcopenia is characterized by progressive loss of skeletal muscle mass, strength, and physical function with aging. Motor unit degeneration is a major contributor, but the mechanisms driving age-related deterioration of motor neurons, neuromuscular junctions (NMJs), and muscle fibers remain incompletely understood. Norepinephrine (NE) regulates spinal motor neuron excitability, but its role in motor unit aging has not been established.

**Methods:** Whole-cell patch-clamp electrophysiology, optogenetics, fast-scan cyclic voltammetry, microdialysis, high-performance liquid chromatography, western blotting, and chemogenetics were used in mice aged 1-25 months. Motor neuron intrinsic properties were examined in 135 lumbar motor neurons from 81 mice. NE release dynamics, tissue content, and expression of the NE transporter (NET) and α2A-adrenergic autoreceptor were assessed in adult and aged mice. Descending spinally projecting noradrenergic neurons were selectively inhibited in aged mice using a retrograde chemogenetic approach.

**Results:** Aging produced progressive motor neuron hyperexcitability characterized by membrane depolarization, increased input resistance, elevated firing frequency, and altered action potential threshold (all P < 0.005). Optogenetically evoked endogenous NE increased motor neuron excitability by reducing action potential latency (P = 0.0001) and decreasing rheobase (P = 0.0028); these effects were prevented by α1-adrenergic receptor blockade (P = 0.0313). Aging enhanced stimulus-evoked spinal NE release, with greater responses at 20 Hz (P = 0.0223), 30 Hz (P = 0.0022), and 40 Hz (P < 0.0001). NE release elicited by stimulation of locus coeruleus and A5 noradrenergic inputs was also increased in aged mice (P = 0.0349 and P = 0.0016, respectively). Basal extracellular NE concentrations were unchanged, whereas spinal cord NE content increased (P = 0.0403). Aging reduced α2A-adrenergic receptor (P = 0.0009) and NET expression (P = 0.0046). Chemogenetic suppression of descending noradrenergic neurons in aged mice increased hindlimb muscle mass (P = 0.0111-0.0344), muscle fiber cross-sectional area (P = 0.0078-0.0467), NMJ innervation (P = 0.0210), and motor neuron soma size (P = 0.0353).

**Conclusions:** Aging promotes spinal noradrenergic overdrive characterized by increased NE storage, enhanced activity-dependent NE release, and impaired autoregulatory control. Excessive noradrenergic signaling contributes to motor neuron hyperexcitability and motor unit deterioration. Suppression of descending noradrenergic activity reverses structural features of motor neuron, NMJ, and skeletal muscle aging, identifying spinal noradrenergic signaling as a potential therapeutic target for sarcopenia.

## Introduction

Sarcopenia is a hallmark of biological aging and is characterized by progressive declines in skeletal muscle mass, strength and physical performance. Although intrinsic alterations within skeletal muscle contribute to this syndrome, accumulating evidence indicates that degeneration of the nervous system is a major determinant of age-related muscle loss. Motor unit remodeling, loss of motor neurons, impaired neuromuscular junction (NMJ) transmission and reduced motor unit recruitment all contribute to the development of sarcopenia.[1–3] Because motor neurons, NMJs and the muscle fibers they innervate function as an integrated motor unit, age-related deterioration of any component can compromise the entire neuromuscular system. Identifying upstream mechanisms that simultaneously influence multiple components of the motor unit is therefore essential for understanding neuromuscular aging and for developing interventions capable of modifying disease progression rather than simply mitigating its consequences.

Descending monoaminergic pathways are critical regulators of spinal motor function. Among these systems, norepinephrine (NE) exerts powerful effects on motor neuron excitability through activation of adrenergic receptors expressed on motor neurons and spinal interneurons. NE enhances persistent inward currents, facilitates repetitive firing and amplifies motor output through α1-adrenergic receptor signaling.[4] While these actions are necessary for normal motor control, sustained enhancement of neuronal excitability may become maladaptive during aging, potentially compromising long-term motor unit stability.

The principal sources of spinal NE arise from the locus coeruleus (LC) and the pontine A5 noradrenergic cell group.[5–7] Age-related degeneration and dysfunction of central noradrenergic pathways have been reported in both rodents and humans and have been linked to cognitive decline, neurodegenerative disease and impaired autonomic regulation.[8–11] However, little is known about how aging affects spinal noradrenergic neurotransmission or whether alterations in NE signaling contribute to degeneration of the motor unit.

NE transmission is tightly regulated by uptake through the NE transporter (NET) and by presynaptic α2-adrenergic autoreceptors that limit transmitter release.[12] Disruption of either mechanism can amplify extracellular NE signaling and enhance neuronal excitation. We therefore hypothesized that aging disrupts spinal noradrenergic homeostasis, producing a state of noradrenergic overdrive characterized by excessive activity-dependent NE signaling and motor neuron hyperexcitability. Such a mechanism could provide a direct link between age-related neurochemical changes in the spinal cord and deterioration of the motor unit.

To test this hypothesis, we examined lumbar motor neuron electrophysiological properties across the lifespan, quantified spinal NE release and metabolism using fast-scan cyclic voltammetry, microdialysis and high-performance liquid chromatography, and assessed age-related changes in NET and α2A-adrenergic receptor expression. We then used chemogenetic inhibition of spinally projecting noradrenergic neurons to determine whether suppressing excessive NE signaling could restore structural features of the aged motor unit. Our findings identify spinal noradrenergic overdrive as a previously unrecognized mechanism of neuromuscular aging and demonstrate that key features of motor unit degeneration remain reversible late in life.

## Methods

### Study design

The study was designed to determine whether aging alters spinal noradrenergic neurotransmission and whether such alterations contribute to motor unit aging. Electrophysiological, neurochemical and molecular approaches were used to characterize age-associated changes in spinal NE signaling. Chemogenetic inhibition of spinally projecting noradrenergic neurons in aged mice was then used to assess whether structural features of motor unit aging, including motor neuron morphology, neuromuscular junction integrity and skeletal muscle fiber size, remained amenable to intervention.

### Animals

C57BL/6 mice aged 1-25 months were obtained from the National Institute on Aging (USA), ChAT-GFP mice (The Jackson Laboratory, strain #007902; Bar Harbor, ME, USA) and TH-Ai32/ChAT-GFP mice aged 2–4 months—generated by crossing TH-Cre1 (JAX #008601), Ai32 (JAX #024109), and ChAT-GFP lines—were used in this study. Equal numbers of male and female mice were included. All mice were bred and housed in a specific pathogen-free facility within the Wake Forest University School of Medicine (WFUSM) Animal Research Program and maintained at 21 °C under a 12:12-h light/dark cycle, with ad libitum access to standard chow and drinking water. For chemogenetic experiments, mice received either plain drinking water or water supplemented with deschloroclozapine (DCZ), as described below.

### Chemogenetics

To selectively inhibit descending noradrenergic neurons projecting to the lumbar spinal cord, a retrogradely transported CAV2-PRS-HA-hM4Di(E1)-hSyn-mCherry(E3) viral vector (Institut de Génétique Moléculaire de Montpellier, University of Montpellier, CNRS, France) was injected into the L4–L5 spinal cord. The virus was diluted to 10⁹ particles in PBS containing 10% glycerol and delivered in a total volume of 500 nL. Eight weeks after injection, mice were assigned to receive either the DREADD agonist deschloroclozapine (DCZ; 10 µg ml⁻¹) in drinking water (n = 10; 5 males, 5 females) or regular drinking water (n = 10; 5 males, 5 females). The PRS promoter restricts hM4Di expression to catecholaminergic neurons, whereas hSyn-driven mCherry permits visualization of all transduced neurons. Hemagglutinin (HA) immunostaining was used to verify DREADD expression in targeted noradrenergic neurons. Four to six weeks later, mice were euthanized, and tissues were collected for analysis, including the lumbar spinal cord, brainstem, and extensor digitorum longus (EDL), soleus, tibialis anterior, and gastrocnemius muscles (Fig. 6B). The EDL and soleus were processed for analyses of myofiber cross-sectional area and myosin heavy-chain expression, while contralateral muscles were used to assess postterminal neuromuscular junction innervation.

### Motor Unit Analysis: Neuromuscular Junction Morphology

EDL and soleus muscles were harvested following transcardial perfusion with saline followed by 4% PFA in 0.2 M sodium phosphate buffer (pH 7.4). Muscles were pinned onto Sylgard-coated dishes, and individual fascicles were carefully dissected and post-fixed in 4% PFA for 20 min. Preparations were washed three times in PBS containing 0.1% Tween-20 (PBST) and blocked overnight at 4°C in 10% goat serum and 1% Triton X-100.

Muscles were incubated overnight at 4°C with antibodies against neurofilament (SMI-312, BioLegend, San Diego, CA, USA) and SV2 (Developmental Studies Hybridoma Bank, University of Iowa, USA) in PBST containing 4% goat serum and 1% Triton X-100 (Supplementary Table 1). Following three washes in PBST, tissues were incubated with Alexa Fluor 488-conjugated goat anti-mouse secondary antibody (1:500) together with tetramethylrhodamine-conjugated α-bungarotoxin (1:250). Neuromuscular junctions (NMJs) were imaged using an Olympus FV4000 confocal microscope with a 60× objective. Z-stack images were reconstructed and analyzed using Fiji/ImageJ (NIH) to quantify NMJ volume, fragmentation, and pre- and postsynaptic colocalization. Briefly, image stacks were thresholded and converted to binary images. NMJ volume was quantified using the 3D Objects Counter plugin, whereas pre- and postsynaptic colocalization was assessed using the Coloc 2 plugin, which calculates pixel-intensity correlation metrics, including Manders’ coefficients, to quantify the spatial overlap of fluorescence signals. Manders’ colocalization coefficients M1 and M2 were used to quantify overlap between presynaptic (NF+SV2) and postsynaptic (α-bungarotoxin) signals. M1 represents the fraction of presynaptic signal overlapping the postsynaptic endplate, whereas M2 represents the fraction of postsynaptic signal covered by the presynaptic terminal. NMJs were further classified according to innervation status based on thresholded Manders coefficient 2 (tM2) values as fully innervated (FI; tM2 ≥ 0.80), early partially denervated (EPD; 0.45 < tM2 < 0.80), advanced partially denervated (APD; 0.10 < tM2 ≤ 0.45), or fully denervated (FD; tM2 ≤ 0.10).

### Motor Unit Analysis: Spinal Motor Neuron Morphology

Following transcardial perfusion with PBS and 4% PFA, lumbar spinal cords were dissected after laminectomy and post-fixed in 4% PFA at 4°C for 24 h. Tissue was cryoprotected overnight at 4°C in 30% sucrose prepared in PBS containing 0.1% Tween-20. Lumbar spinal cord segments were embedded in O.C.T. compound and coronally sectioned from L3–L5 at a thickness of 10 μm using a Leica CM3050S cryostat.

Sections were blocked overnight at 4°C in 10% donkey serum and 1% Triton X-100 and then incubated overnight at 4°C with anti-choline acetyltransferase (ChAT) antibody (MilliporeSigma; Supplementary Table 1). The following day, sections were incubated with Alexa Fluor 488-conjugated donkey anti-goat secondary antibody (Invitrogen, Thermo Fisher Scientific). Nuclei were counterstained with Hoechst 33342 (Thermo Fisher Scientific) for 5 min at room temperature.

Sections were mounted with fluorescence mounting medium (Dako) and imaged using an Olympus FV4000 confocal microscope. Maximum-intensity projections were analyzed using Fiji/ImageJ for background subtraction, thresholding, segmentation, and quantification of motor neuron number and soma area. To avoid double-counting, every other section was analyzed, and only motor neurons containing a clearly identifiable nucleus were included in the quantification. Analyses were performed within ventral horn regions corresponding to lumbar motor neuron pools that innervate the hindlimb muscles examined in this study.

Sources, catalog numbers, and concentrations of all pharmacological agents used in this study are listed in Supplementary Table 2.

### Data Analysis

Data were analyzed using GraphPad Prism 11 (Dotmatics, Boston, MA, USA) and Microsoft Excel (Microsoft Corporation, Redmond, WA, USA). Statistical tests used for each experiment are indicated in the corresponding figure legends. Equal numbers of male and female mice were included in all experimental groups. Preliminary analyses revealed no consistent sex-dependent trends across the primary outcome measures, and data from both sexes were therefore combined to maximize statistical power. The study was not specifically powered to evaluate sex effects. Data are presented as mean ± s.d. unless otherwise stated. Individual data points represent biological replicates unless noted otherwise. Statistical significance was defined as *P* < 0.05. Statistical tests, exact *P* values, and sample sizes for all analyses are provided in Supplementary Table 3.

For patch-clamp electrophysiology, skeletal muscle immunohistochemistry, fast-scan voltammetry, microdialysis, norepinephrine (NE) content measurements, and immunoblotting experiments, please refer to the *Supplementary Methods*.

## Results

### Aging promotes motor neuron hyperexcitability

Whole-cell patch-clamp recordings from L4–L5 spinal motor neurons in mice aged 1–25 months revealed marked age-dependent changes in intrinsic properties (Fig. 1e–i). α-motor neurons were identified based on large soma size and characteristic morphology, location within Rexed laminae VIII and IX, and the absence of spontaneous firing. In contrast, γ-motor neurons exhibit spontaneous activity,[13] whereas glial cells do not generate action potentials[14]. Motor neuron identity was further confirmed post-hoc by choline acetyltransferase (ChAT) immunostaining (Fig. 1j-m). Resting membrane potential became progressively depolarized with age (P < 0.0001), while membrane capacitance declined significantly (P < 0.0001), consistent with reduced somatic size. Input resistance increased (P < 0.0001), amplifying responsiveness to synaptic inputs. Cumulative firing frequency rose sharply (P < 0.0001), and action potential threshold shifted significantly with age (P < 0.005), indicating altered voltage-gated channel dynamics. These changes collectively point to age-related motor neuron hyperexcitability. These findings indicate that aging progressively shifts lumbar motor neurons toward a hyperexcitable state that may increase vulnerability of the motor unit to long-term dysfunction and degeneration. Exact sample sizes, statistical tests, and *P* values for all analyses are summarized in Supplementary Table 3.

**Figure 1.**
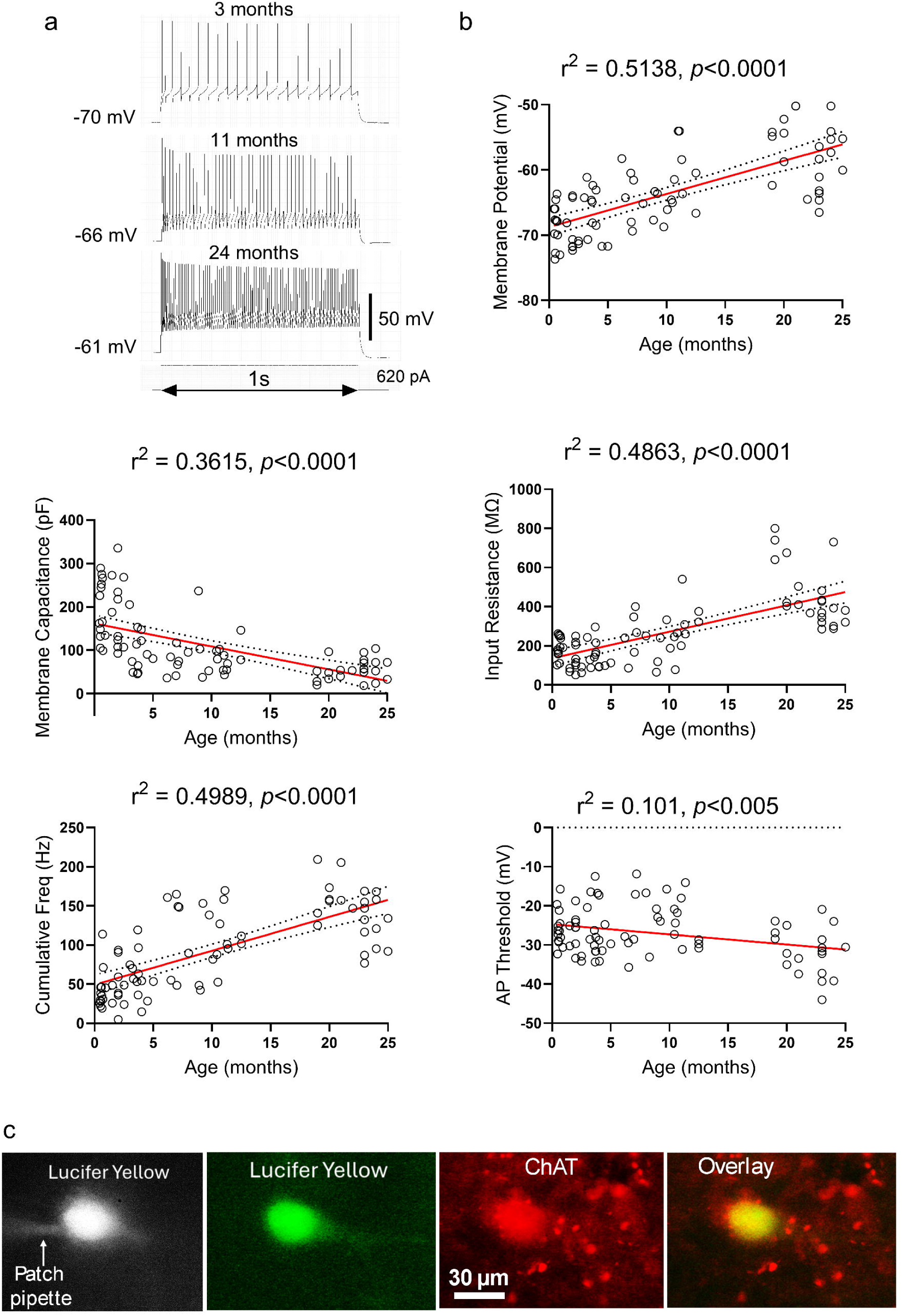
Aging is associated with progressive motor neuron hyperexcitability and structural remodeling. (a) Representative whole-cell current-clamp recordings obtained from lumbar spinal motor neurons of 3-, 11-, and 24-month-old mice during a 620-pA depolarizing current injection. Aging was associated with progressive membrane depolarization and increased firing frequency. (b) Relationships between age and motor neuron intrinsic electrophysiological properties. Aging was associated with progressive membrane depolarization (top right; *r*² = 0.5138, *p* < 0.0001), reduced membrane capacitance (middle left; *r*² = 0.3615, *p* < 0.0001), increased input resistance (middle right; *r*² = 0.4863, *p* < 0.0001), increased cumulative firing frequency during depolarizing current injection (bottom left; *r*² = 0.4989, *p* < 0.0001), and a modest reduction in action potential threshold (bottom right; *r*² = 0.101, *p* < 0.005). Each symbol represents one motor neuron. Red lines indicate linear regression fits; dotted lines indicate 95% confidence intervals. (c) Post hoc identification of a patch-clamped motor neuron filled with Lucifer Yellow. Images show Lucifer Yellow fluorescence, choline acetyltransferase (ChAT) immunoreactivity, and the merged image confirming that the recorded neuron was ChAT-positive. Scale bar = 30 μm. Data were obtained from 135 lumbar motor neurons recorded from 81 mice aged 1-25 months. Values in scatter plots represent individual motor neurons.

### Noradrenergic activation drives motor neuron depolarization

To determine the impact of NE signaling on motor neuron excitability, we optogenetically stimulated NE neurons in young TH-Ai32/ChR2-GFP × ChAT-GFP mice while recording lumbar motor neurons. Optostimulation produced repeated bursts of action potentials (APs) and sustained membrane depolarizations (Fig. 2a), accompanied by an increase in excitatory postsynaptic potentials (EPSPs) compared with baseline (Fig. 2b). Using a 400-ms ramp protocol from −70 mV to −45 mV to assess excitability near firing threshold, we observed that NE release significantly advanced the onset of AP generation and increased firing frequency (Fig. 2c-f), with responses returning to baseline within ∼2 min. Quantitative analysis revealed a marked reduction in AP latency following optostimulation (*P* = 0.0001; Fig. 2g) and a significant decrease in the current required to elicit APs (*P* = 0.0028; Fig. 2h).

**Figure 2.**
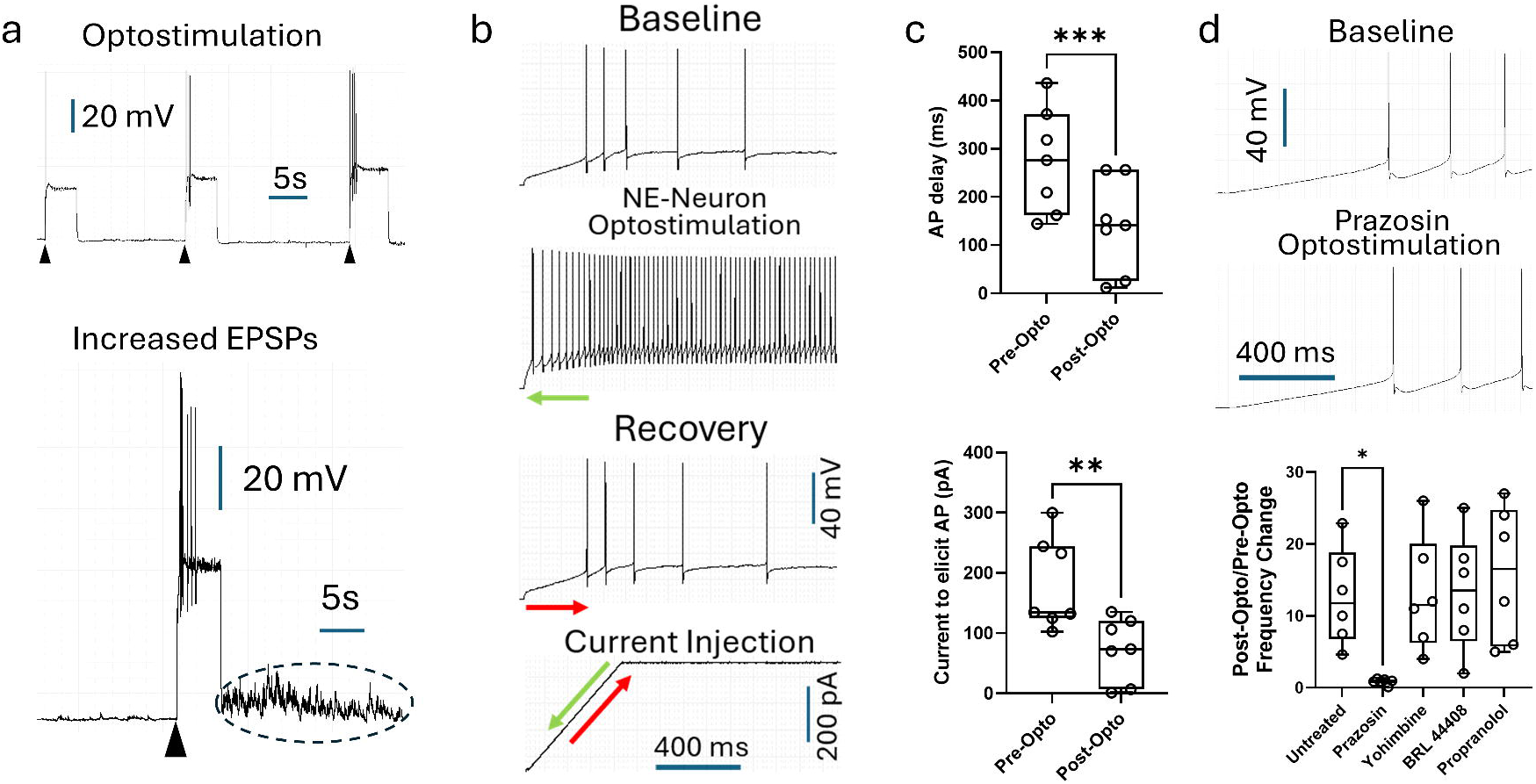
Optogenetic activation of noradrenergic neurons enhances motor neuron excitability through α1-adrenergic receptor signaling. (a) Representative whole-cell recordings from lumbar motor neurons in young (2-4-month-old) TH-Ai32/ChR2-GFP × ChAT-GFP mice during optogenetic stimulation of noradrenergic neurons (arrowheads). Optostimulation evoked bursts of action potentials and prolonged membrane depolarization (top) and increased excitatory postsynaptic potential (EPSP) activity following norepinephrine release (bottom, dashed ellipse). (b) Representative recordings illustrating action potential generation during a 400-ms depolarizing current ramp. Traces obtained under baseline conditions (top), immediately after noradrenergic neuron optostimulation (middle), and following recovery (∼2 min; bottom) demonstrate earlier action potential initiation and enhanced excitability after endogenous norepinephrine release. The schematic below shows the ramp current injection protocol. Green arrows indicate advanced action potential onset after optostimulation, and red arrows indicate recovery toward baseline. (c) Quantification of motor neuron excitability before and after optostimulation. Optogenetically evoked norepinephrine release significantly reduced action potential latency (top) and decreased the amount of injected current required to elicit action potentials (bottom), indicating increased motor neuron excitability. (d) Pharmacological characterization of the adrenergic receptor subtype mediating the excitatory effect of endogenous norepinephrine. Representative traces obtained before (top) and after incubation with the α1-adrenergic receptor antagonist prazosin (10 μM, 5 min; bottom) demonstrate attenuation of the excitatory response to optostimulation. Quantification of firing frequency changes following optostimulation in untreated preparations and in the presence of adrenergic receptor antagonists showed that only prazosin significantly reduced the response, whereas the α2-adrenergic receptor antagonist yohimbine, the β1-adrenergic receptor antagonist BRL 44408, and the β-adrenergic receptor antagonist propranolol had no significant effect. Data are presented as mean ± SEM. Box plots show the median, interquartile range, and individual data points. Statistical significance is indicated as *P* < 0.05, P < 0.01, and *P* < 0.001.

To identify the adrenergic receptor subtype mediating this response, motor neurons were exposed to adrenergic receptor antagonists before optostimulation. Pretreatment with the α1-adrenergic receptor (α1-AR) antagonist prazosin (10 μM, 5 min) abolished the excitatory effects of optostimulation, reducing the post-/pre-optostimulation firing-frequency ratio from 12.69 to 0.87 (adjusted *P* = 0.0313; Fig. 2i-k). In contrast, neither the α2-adrenergic receptor (α2-AR) antagonists yohimbine (10 μM; adjusted *P* > 0.9999) and BRL 44408 (2 μM; adjusted *P* = 0.9996), nor the β-adrenergic receptor (β-AR) antagonist propranolol (10 μM; adjusted *P* = 0.8660), altered the response to optostimulation (Fig. 2k). One-way ANOVA followed by Dunnett’s multiple-comparisons test demonstrated that only α1-AR blockade significantly attenuated the effect of endogenous NE release. These findings indicate that the facilitation of motor neuron excitability by endogenous NE is mediated primarily through α1-adrenergic receptor signaling.

Although optogenetic stimulation demonstrated that catecholamine release enhances motor neuron excitability, the identity of the transmitter responsible and the mechanisms regulating its availability in the spinal cord remained unclear. We therefore used fast-scan cyclic voltammetry together with pharmacological manipulations of monoamine storage, release, uptake and autoreceptor signaling to define the catecholamine detected in lumbar spinal cord slices and to identify pathways controlling its extracellular concentration.

### Catecholamine release is regulated by transporters and α2 autoreceptors

Fast-scan cyclic voltammetry in lumbar spinal slices revealed robust catecholamine release upon electrical stimulation (Fig. 3a,b). Cocaine significantly enhanced release (P < 0.05), whereas reserpine markedly reduced it (P < 0.01) (Fig. 3c,d). Net blockade with desipramine, GBR12909 and fluoxetine had no effect (Fig. 3e,h,i), supporting NE as the primary catecholamine detected. α2 antagonism (idazoxan) increased release (P < 0.05), whereas α2 agonism (clonidine) decreased it (P < 0.05) (Fig. 3j,k), consistent with presynaptic α2 autoreceptor feedback inhibition. Dopaminergic agents, including the D2 receptor agonist quinpirole and antagonist raclopride, had no effect on evoked catecholamine release, further supporting norepinephrine as the predominant catecholamine detected (Fig. 3f,g; Supplementary Figs. 2–3).

**Figure 3.**
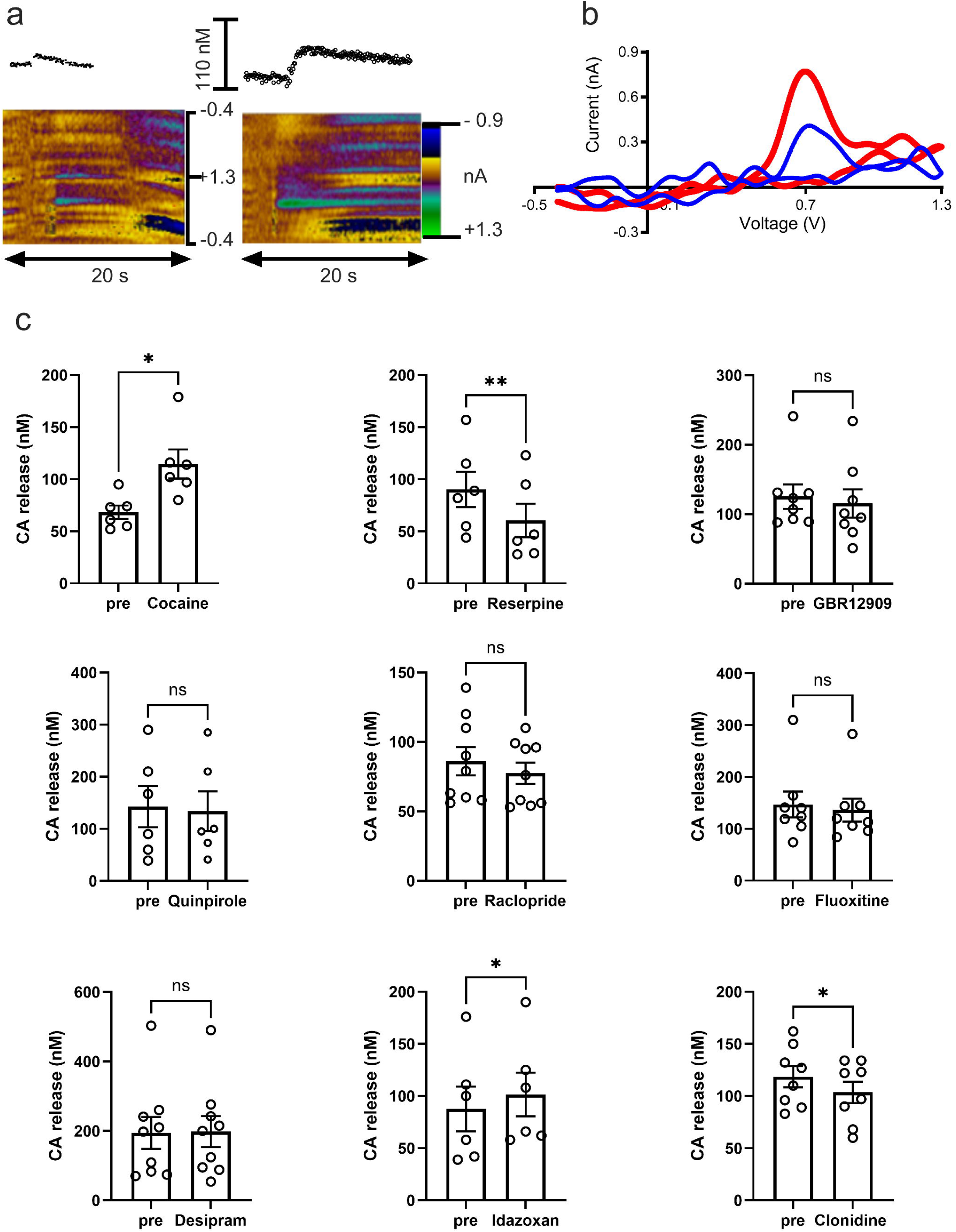
Pharmacological characterization of electrically evoked catecholamine release in lumbar spinal cord slices. (a) Representative fast-scan cyclic voltammetry recordings demonstrating electrically evoked catecholamine release before and after inhibition of monoamine reuptake with cocaine (10 μM). Two-dimensional color plots show electrochemical signals as a function of time (x-axis), applied potential (y-axis), and background-subtracted faradaic current (pseudo-color scale). Traces above each color plot illustrate changes in extracellular catecholamine concentration, demonstrating that cocaine increases the amplitude of the evoked signal. (b) Representative cyclic voltammograms obtained before (blue) and after (red) cocaine application. The characteristic oxidation and reduction currents confirm the electrochemical detection of catecholamines and demonstrate enhancement of the evoked signal following cocaine treatment. (c) Summary of pharmacological manipulations affecting electrically evoked catecholamine release. Inhibition of monoamine transporters with cocaine (10 μM) significantly increased evoked catecholamine release, whereas depletion of vesicular catecholamine stores with reserpine (10 μM) significantly reduced the response. Blockade of dopamine transporters with GBR12909 (5 μM), activation of dopamine D2 receptors with quinpirole (10 μM), inhibition of dopamine D2 receptors with raclopride (10 μM), inhibition of serotonin transporters with fluoxetine (10 μM), and inhibition of norepinephrine transporters with desipramine (10 μM) did not significantly alter evoked catecholamine release. In contrast, blockade of α2-adrenergic receptors with idazoxan (10 μM) significantly enhanced evoked release, whereas activation of α2-adrenergic receptors with clonidine (10 μM) significantly reduced the response. These findings identify norepinephrine as the predominant catecholamine signal detected in the lumbar spinal cord and demonstrate regulation of release through α2-adrenergic autoreceptors. Bars represent mean ± SEM and symbols represent individual mice. Statistical comparisons were performed using paired *t*-tests. *P* < 0.05, P < 0.01; ns, not significant.

### Aging enhances activity-dependent spinal norepinephrine release

We next examined whether aging alters NE release dynamics. Electrical stimulation elicited frequency-dependent NE efflux in both adult and old mice, with significantly greater responses at 20–40 Hz in aged animals (P < 0.01 at 30 Hz; P < 0.0001 at 40 Hz) (Fig. 4a). In vivo stimulation of the LC (Fig. 4b) and the ventrolateral pontine region containing A5 noradrenergic neurons (Fig. 4c,d) evoked significantly greater NE release in aged mice (LC: P<0.05, A5: P < 0.01). Stimulation of the LC produced robust local NE release but failed to evoke detectable release in the A5 region. Conversely, stimulation of the A5 region produced robust local NE release without detectable release in the LC. These findings indicate that LC and A5 noradrenergic pathways operate independently under the present experimental conditions. Supplementary Fig. 4 illustrates stimulation electrode placement adjacent to pontine A5 neurons, while LC electrode positioning has been reported previously.[15] Collectively, these findings demonstrate that the aged spinal cord is exposed to amplified NE transients during physiological activation of descending noradrenergic pathways, creating conditions that favor excessive motor neuron excitation.

**Figure 4.**
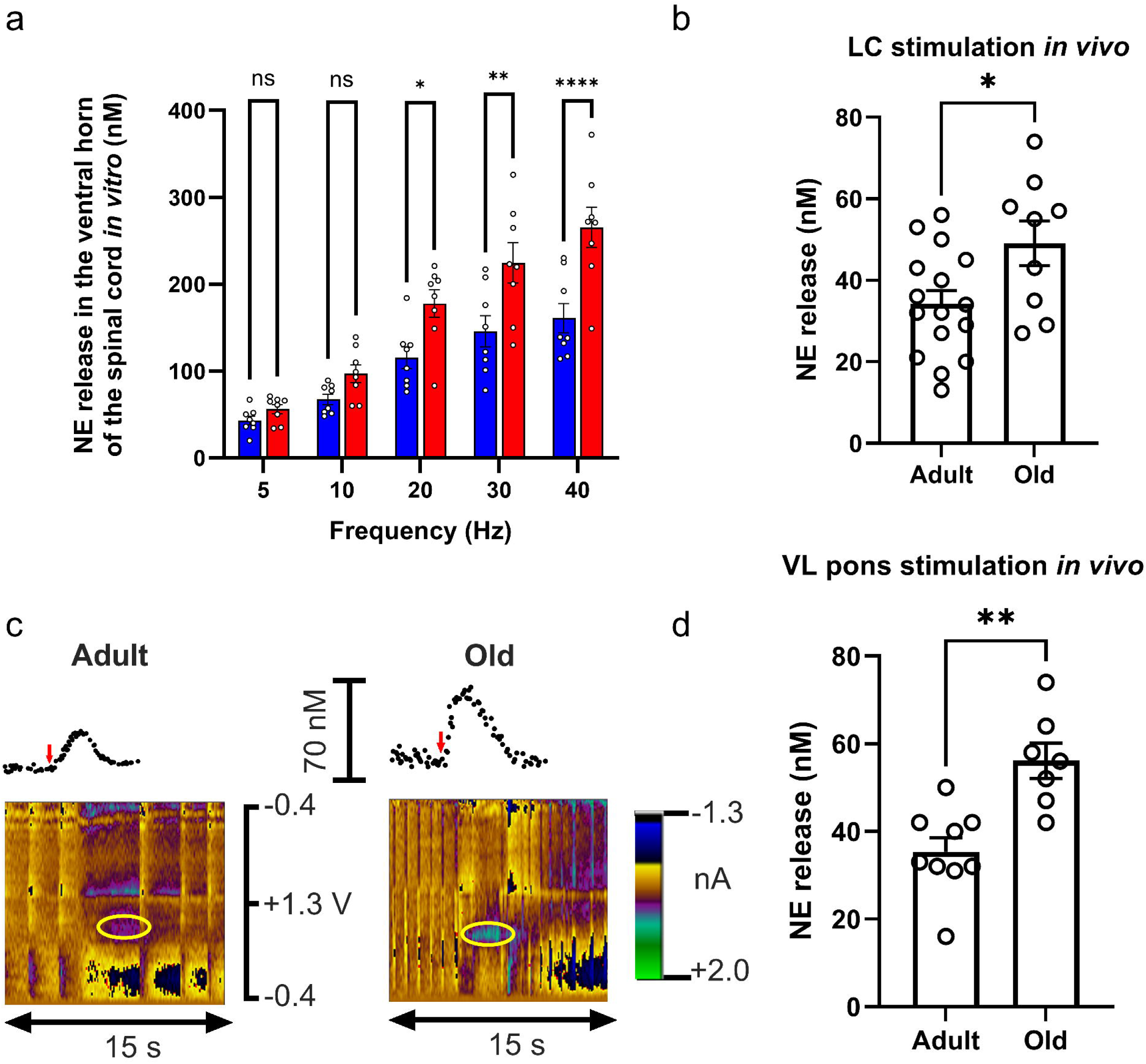
Aging amplifies norepinephrine release in lumbar spinal cord in vitro and in vivo. (a) Electrical stimulation (5–40 Hz, 1 s) elicited frequency-dependent NE efflux in adult (6–7 months) and aged (22–23 months) mice, with significantly greater responses at 20–40 Hz in aged animals. Bars show mean ± SEM (n = 8 per group). Two-way ANOVA with Sidak’s multiple comparisons: *P < 0.05, **P < 0.01, ****P < 0.0001; ns, not significant. b NE concentration (nM) measured in ventral horn after LC stimulation at 40 Hz was significantly higher in aged mice (Unpaired t-test, P < 0.05). (c-d) NE release following ventrolateral pons stimulation was significantly greater in aged mice (Unpaired t-test, P < 0.01). Individual points represent biological replicates.

### Aging increases spinal norepinephrine stores while preserving basal extracellular levels

Microdialysis revealed no significant difference in extracellular NE or MHPG between adult and old mice (Fig. 5a,b). However, tissue analysis showed a significant increase in NE content in aged spinal cords (P = 0.0403) (Fig. 5c), while MHPG tended to decrease (Fig. 5d). Neuromelanin and dopamine levels remained unchanged (Fig. 5e,f). These findings suggest that aging increases NE storage capacity without elevating basal extracellular levels, thereby preserving resting neurotransmitter concentrations while exaggerating activity-dependent signaling. Such enhanced phasic NE release would be expected to disproportionately affect motor neurons during periods of sustained activation.

**Figure 5.**
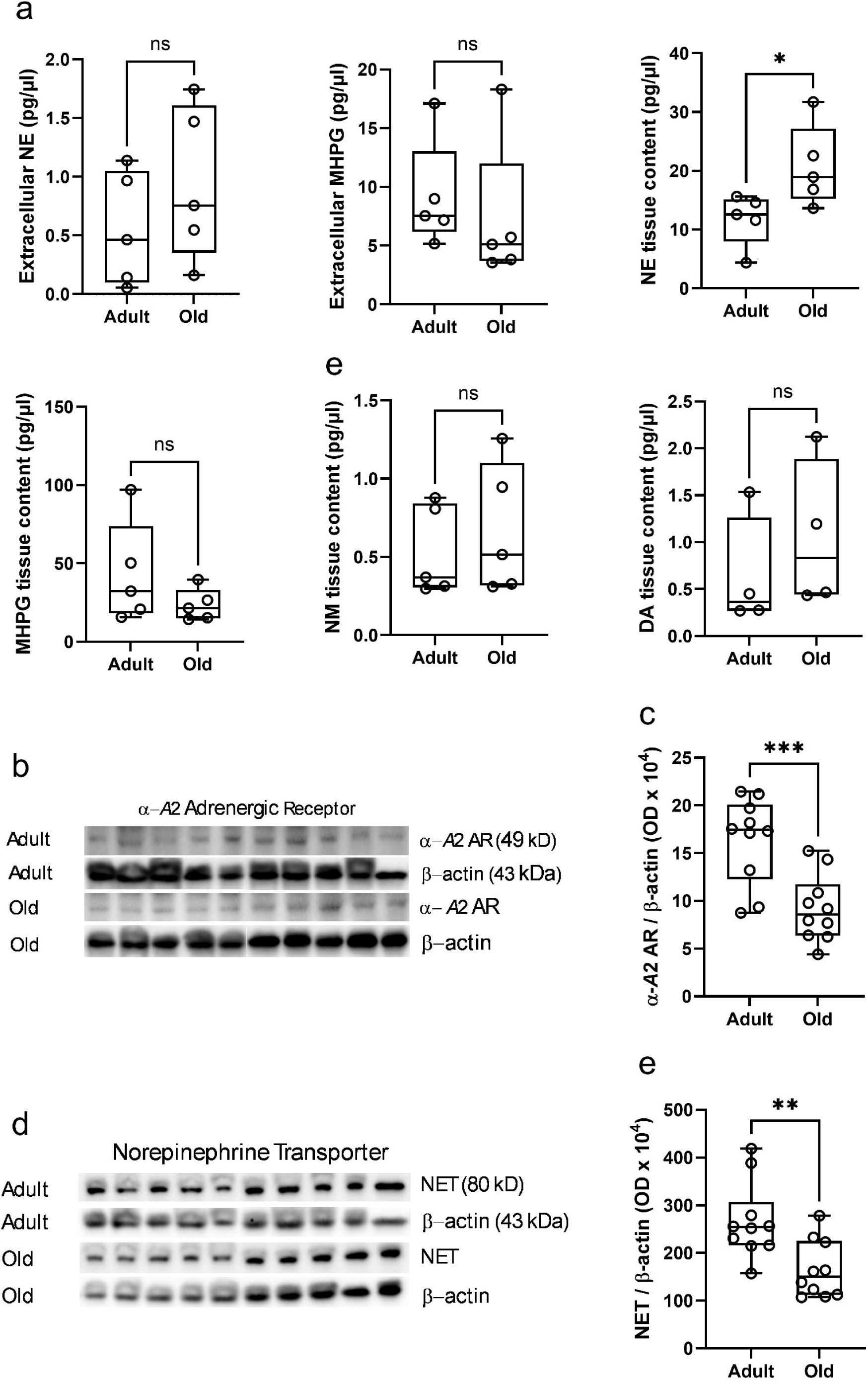
Aging increases spinal norepinephrine content and reduces expression of key regulators of noradrenergic signaling. (a) Neurochemical analysis of lumbar spinal cord tissue from adult and aged mice. Extracellular norepinephrine (NE) and its principal metabolite, 3-methoxy-4-hydroxyphenylglycol (MHPG), were measured by in vivo microdialysis. Tissue concentrations of NE, MHPG, neuromelanin (NM), and dopamine (DA) were quantified by high-performance liquid chromatography with electrochemical detection. Basal extracellular NE and MHPG concentrations were not significantly different between age groups. In contrast, spinal cord tissue NE content was significantly elevated in aged mice (*P* = 0.0403), whereas tissue MHPG, NM, and DA levels were unchanged. (b) Representative immunoblots of α2A-adrenergic receptor (α2A AR; 49 kDa) and β-actin (43 kDa) expression in lumbar spinal cord homogenates obtained from adult and aged mice. (c) Quantification of α2A AR protein expression normalized to β-actin optical density. Aged mice exhibited a significant reduction in α2A AR expression compared with adult mice (*P* = 0.0009). (d) Representative immunoblots of norepinephrine transporter (NET; 80 kDa) and β-actin (43 kDa) expression in lumbar spinal cord homogenates from adult and aged mice. (e) Quantification of NET protein expression normalized to β-actin optical density. NET expression was significantly reduced in aged mice compared with adults (*P* = 0.0046). Collectively, these findings indicate that aging is associated with increased spinal NE accumulation despite unchanged basal extracellular NE concentrations, accompanied by reduced expression of the α2A-adrenergic autoreceptor and the NE transporter, two key mechanisms regulating noradrenergic feedback inhibition and neurotransmitter clearance. Box plots show the median, interquartile range, and individual data points (one point per mouse). Statistical comparisons were performed using two-tailed unpaired *t*-tests. Neurochemical concentrations are expressed as pg/μl. *P* < 0.05, *P* < 0.001; ns, not significant.

### Aging impairs autoregulatory control of spinal norepinephrine signaling

Western blot analysis revealed significant reductions in α2A-adrenergic receptor (P < 0.001) and NE transporter (P < 0.01) expression in aged spinal cords (Fig. 5g-j). These changes likely impair presynaptic feedback inhibition and NE clearance, promoting a state of noradrenergic overdrive capable of continuously challenging motor unit homeostasis during aging. Together, these results demonstrate that aging promotes a noradrenergic imbalance characterized by increased NE storage, exaggerated phasic release, and diminished autoreceptor and transporter function. This neurochemical environment drives motor neuron hyperexcitability, providing a mechanistic link to age-related neuromuscular decline.

Together, these findings support a model in which aging produces spinal noradrenergic overdrive characterized by increased NE storage, exaggerated stimulus-evoked release and impaired neurotransmitter regulation. Because excessive NE signaling promotes motor neuron excitability and may compromise motor unit homeostasis, we next tested whether suppressing descending noradrenergic activity could reverse established features of motor unit aging in advanced age.

### Chemogenetic inhibition of spinally projecting noradrenergic neurons improves muscle morphology in aged mice

To determine whether excessive noradrenergic signaling contributes directly to neuromuscular decline, we selectively inhibited spinally projecting noradrenergic neurons using a retrogradely transported CAV2 PRS HA-hM4D (E1) hSyn mCherry (E3) construct injected into the L4–L5 spinal cord (Fig. 6a,b). Viral targeting was confirmed by HA and mCherry expression within tyrosine hydroxylase-positive neurons of the ventrolateral pontine A5 noradrenergic region. Retrograde labeling of A5 neurons following spinal injection was confirmed in Supplementary Fig. 5.

**Figure 6.**
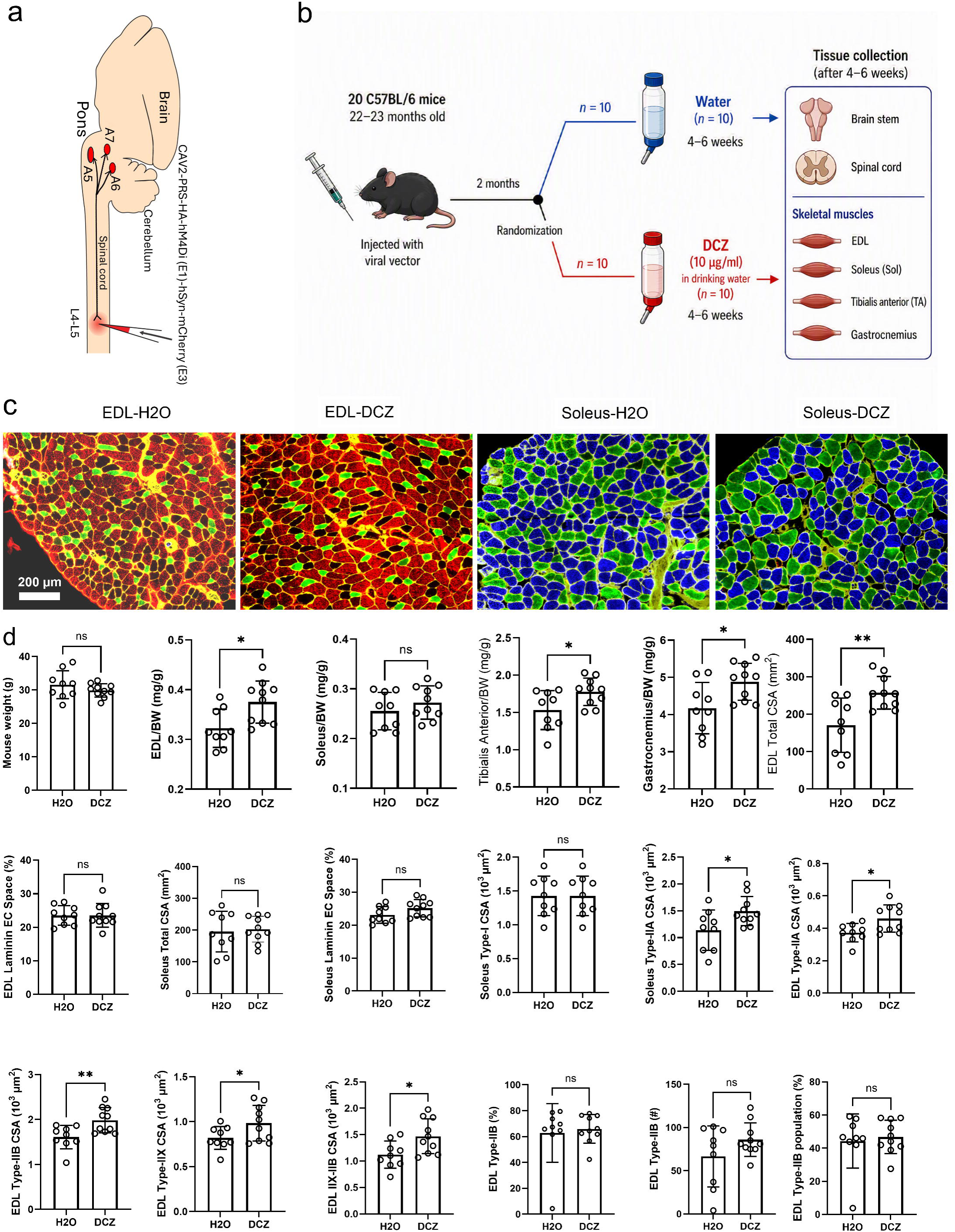
Chemogenetic inhibition of spinally projecting noradrenergic neurons improves skeletal muscle morphology in aged mice. (a) Schematic illustrating bilateral lumbar spinal cord (L4-L5) injections of CAV2-PRS-HA-hM4Di(E1)-hSyn-mCherry(3) and the anatomical distribution of spinally projecting noradrenergic neurons within the pontine A5-A7 nuclei. (b) Experimental design. Aged C57BL/6 mice (22-23 months old) received lumbar spinal cord injections of the retrograde inhibitory DREADD vector. Following a two-month expression period, mice were treated with either drinking water (H₂O; *n* = 10) or deschloroclozapine (DCZ; 10 μg mL⁻¹ in drinking water; *n* = 10) for 4-6 weeks before tissue collection. Brainstem, spinal cord, and hindlimb muscles were harvested for subsequent analyses. (c) Representative laminin-stained cross-sections of extensor digitorum longus (EDL) and soleus muscles from H₂O- and DCZ-treated mice. DCZ-mediated inhibition of descending noradrenergic neurons increased muscle size and reduced extracellular space. Scale bar = 200 μm. (d) Effects of chemogenetic inhibition of spinally projecting noradrenergic neurons on body weight, hindlimb muscle mass, and muscle morphology. Body weight was unchanged between groups. In contrast, DCZ treatment significantly increased EDL, tibialis anterior, and gastrocnemius muscle weights normalized to body weight, whereas soleus muscle weight was not significantly altered. Total EDL cross-sectional area and soleus type I fiber cross-sectional area were increased following DCZ treatment, while extracellular space within both EDL and soleus muscles was reduced. (e) Fiber type-specific analysis of muscle remodeling. Chemogenetic inhibition of descending noradrenergic neurons increased the cross-sectional area of EDL type IIA, IIB, and IIX fibers, as well as the combined type I/IIX fiber population. The abundance, number, and relative proportion of EDL type IIB fibers were also altered, indicating substantial remodeling of fast-twitch muscle fibers in response to reduced noradrenergic drive. These findings demonstrate that chronic suppression of spinally projecting noradrenergic neurons partially reverses age-related skeletal muscle atrophy and promotes remodeling of multiple muscle fiber populations in aged mice. Bars represent mean ± SD, and circles represent individual mice. Statistical comparisons were performed using two-tailed unpaired *t*-tests. *P* < 0.05, P < 0.01; ns, not significant.

Body weight did not differ between H2O- and DCZ-treated animals (Fig. 6g). However, DCZ treatment significantly increased EDL, tibialis anterior and gastrocnemius muscle weight normalized to body weight, whereas soleus mass was unchanged (Fig. 6h–k). Total EDL cross-sectional area was also significantly larger in DCZ-treated mice (Fig. 6l). Analysis of extracellular space revealed no significant differences in either EDL or soleus muscles (Fig. 6m,o), suggesting that the increase in muscle size was not attributable to edema or fibrosis.

Fiber-type analysis demonstrated selective enlargement of fast-twitch fibers. DCZ treatment increased the cross-sectional area of EDL type-IIA, type-IIB and type-IIX fibers (Fig. 6r–t), whereas soleus type-I fibers and total soleus cross-sectional area were unaffected (Fig. 6n,p). Several infrequent fiber populations also remained unchanged (Supplementary Fig. 6). Together, these findings indicate that suppression of noradrenergic activity selectively improves features of aged fast motor units, which are particularly susceptible to degeneration during aging. The increase in fast-fiber size further suggests that structural deterioration of aged motor units remains amenable to intervention even late in life, supporting the concept that key components of sarcopenia retain substantial plasticity.

### Chemogenetic suppression of noradrenergic activity improves EDL neuromuscular junction innervation and motor neuron morphology

Because aging is associated with progressive NMJ denervation, we next examined whether reducing noradrenergic tone affected NMJ integrity. In the EDL, NMJ volume and fragmentation were unchanged by treatment (Fig. 7g,h). However, colocalization between presynaptic and postsynaptic structures, measured by Manders’ M1 coefficient, was significantly increased following DCZ administration (Fig. 7i), indicating improved synaptic apposition. Consistent with this observation, the fraction of fully denervated NMJs was significantly reduced in DCZ-treated animals (Fig. 7k). In contrast, soleus NMJ morphology and denervation status were unaffected (Fig. 7r–v).

**Figure 7.**
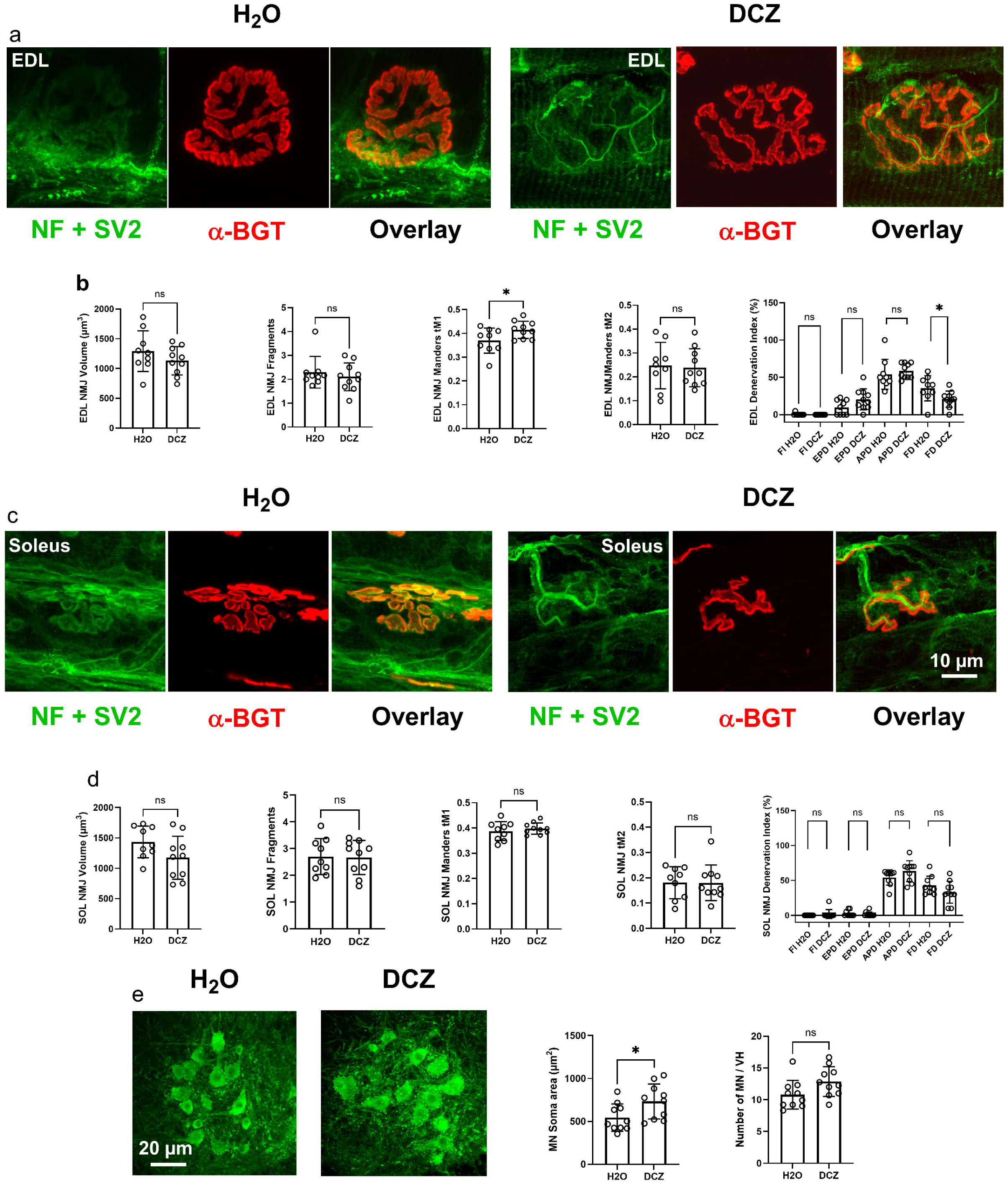
Chemogenetic inhibition of spinally projecting noradrenergic neurons improves neuromuscular junction innervation and motor neuron morphology in aged mice. (a) Representative images of extensor digitorum longus (EDL) neuromuscular junctions (NMJs) from aged mice receiving drinking water (H₂O) or deschloroclozapine (DCZ). Presynaptic terminals were labeled with neurofilament and synaptic vesicle protein 2 (NF + SV2, green), and postsynaptic acetylcholine receptors were labeled with α-bungarotoxin (α-BGT, red). Merged images demonstrate the degree of presynaptic-postsynaptic overlap. (b) Quantification of EDL NMJ morphology. NMJ volume, number of NMJ fragments, Manders’ colocalization coefficients (M1 and M2), and denervation status were assessed. DCZ treatment significantly increased presynaptic-postsynaptic colocalization (M1) and reduced the proportion of fully denervated NMJs, indicating improved NMJ integrity. NMJ volume, fragmentation, and M2 values were not significantly affected. (c) Representative images of soleus NMJs from H₂O- and DCZ-treated mice stained for NF + SV2 (green) and α-BGT (red). Merged images illustrate the relationship between presynaptic terminals and postsynaptic endplates. Scale bar = 10 μm. (d) Quantification of soleus NMJ morphology, including NMJ volume, number of fragments, Manders’ colocalization coefficients (M1 and M2), and denervation categories. No significant differences were detected between treatment groups, indicating that inhibition of descending noradrenergic neurons preferentially affected NMJs in the predominantly fast-twitch EDL muscle. (e) Representative images and quantification of lumbar spinal motor neurons from H₂O- and DCZ-treated mice. Chemogenetic suppression of spinally projecting noradrenergic neurons significantly increased motor neuron soma area without altering the number of motor neurons per ventral horn. Scale bar = 20 μm. These findings demonstrate that reducing descending noradrenergic drive in aged mice partially reverses age-associated motor unit pathology by improving NMJ innervation and increasing motor neuron size, with the most pronounced effects observed in fast-twitch EDL motor units. Bars represent mean ± SD and circles represent individual mice. Denervation index data were analyzed using a mixed-effects model followed by Sidak’s multiple-comparisons test; all other comparisons were performed using two-tailed unpaired *t*-tests with Welch’s correction. *P* < 0.05; ns, not significant.

We also assessed lumbar spinal motor neuron morphology. Chemogenetic inhibition significantly increased motor neuron soma area without affecting motor neuron number per ventral horn (Fig. 7y,z). These findings suggest that excessive noradrenergic activity contributes to motor neuron structural atrophy rather than cell loss and that reducing noradrenergic drive promotes motor neuron maintenance in advanced age.

Collectively, these results demonstrate that chronic suppression of spinally projecting noradrenergic neurons improves multiple structural components of the aged motor unit, including motor neuron morphology, neuromuscular junction innervation and skeletal muscle fiber size. These observations support the concept that age-related noradrenergic dysregulation is not merely associated with motor unit decline but contributes directly to its progression. Importantly, the ability of noradrenergic suppression to ameliorate abnormalities across all three components of the motor unit demonstrates that structural manifestations of neuromuscular aging remain at least partially reversible.

## Discussion

The present study identifies spinal noradrenergic overdrive as a previously unrecognized and reversible mechanism of motor unit aging. We show that aging disrupts multiple components of spinal NE homeostasis, resulting in amplified activity-dependent NE release, motor neuron hyperexcitability and progressive deterioration of motor unit integrity. Importantly, suppressing descending noradrenergic activity in aged mice improved motor neuron morphology, neuromuscular junction innervation and skeletal muscle structure, demonstrating that key manifestations of neuromuscular aging remain amenable to intervention. Together, these findings establish spinal NE signaling as an upstream regulator of motor unit decline and a potential therapeutic target for sarcopenia.

Previous work has emphasized the importance of reduced neuromuscular connectivity and motor unit remodeling in sarcopenia.[1, 16] Our findings suggest that these processes may be driven, in part, by chronic overactivation of spinal motor circuits.[17] Although NE is essential for normal movement and motor neuron recruitment, sustained increases in excitability may impose a long-term metabolic and electrophysiological burden on aging neurons. Our optogenetic and pharmacological studies demonstrate that endogenous NE increases motor neuron excitability primarily through α1-adrenergic receptor signaling, advancing spike initiation and reducing the depolarizing drive required for motor neuron recruitment. Similar forms of neuronal hyperexcitability have been implicated in amyotrophic lateral sclerosis and other neurodegenerative disorders, where excessive neuronal activity accelerates structural deterioration and synaptic loss.[18–20] These findings suggest that chronic activation of α1-adrenergic signaling may contribute to the age-associated loss of motor unit integrity observed in the present study.

A notable aspect of the present findings is that a single neuromodulatory system appears capable of influencing all major components of the motor unit. Aging-related changes in motor neurons, neuromuscular junctions and muscle fibers are often studied independently. However, our results support a model in which excessive spinal NE signaling contributes to coordinated deterioration across the entire motor unit. The observation that suppression of noradrenergic activity improved structural features at each of these levels suggests that spinal noradrenergic overdrive functions as an upstream regulator of neuromuscular aging rather than a consequence of degeneration occurring elsewhere in the system.

Several observations support the existence of noradrenergic overdrive in the aged spinal cord. First, tissue NE content was elevated despite unchanged basal extracellular concentrations. Second, electrically evoked NE release was markedly enhanced in aged animals. Third, expression of both the α2A autoreceptor and NET was reduced. Together, these findings suggest a model in which noradrenergic terminals accumulate larger stores of NE while simultaneously losing critical mechanisms that limit release and promote transmitter clearance. Under resting conditions extracellular NE may remain relatively stable, but physiologically relevant activation of descending pathways produces disproportionately large NE transients. Given our finding that endogenous NE facilitates motor neuron excitation predominantly through α1-adrenergic receptors, these amplified NE signals are likely to enhance motor neuron recruitment and firing during normal motor activity, thereby providing a mechanistic link between spinal noradrenergic overdrive and age-related motor neuron hyperexcitability.

The involvement of both LC and A5 noradrenergic neurons is particularly noteworthy. Descending projections from these nuclei are known to regulate motor output, autonomic function and nociceptive processing.[21–23] Our data indicate that aging alters the functional consequences of activation within both pathways and that exaggerated NE release converges on the lumbar motor network. The independence of LC and A5 signaling demonstrated by FSCV recordings suggests that multiple noradrenergic sources contribute in parallel to spinal NE overdrive.

The chemogenetic experiments provide important causal evidence. Suppression of spinally projecting noradrenergic neurons increased fast-muscle fiber size, reduced EDL denervation and enlarged motor neuron somata. These effects were most pronounced in fast motor units, which are preferentially lost during aging and are strongly associated with declines in strength and mobility.[3, 24, 25] The improvement in NMJ integrity further supports the concept that excessive noradrenergic signaling contributes to failure of motor unit maintenance. Importantly, these improvements were observed in mice that were already 22–23 months old at the onset of treatment, indicating that major structural features of motor unit aging remain modifiable late in life. This degree of reversibility is particularly encouraging because it suggests that neuromuscular aging is not exclusively driven by irreversible neuronal loss.

From a translational perspective, these findings are particularly important because intervention was initiated after substantial aging had already occurred. Suppression of descending noradrenergic activity in advanced age improved structural features of motor neurons, neuromuscular junctions and skeletal muscle fibers, indicating that multiple components of the motor unit retain substantial plasticity late in life. These observations suggest that age-related motor decline is not simply the consequence of cumulative and irreversible tissue loss, but is also driven by an active neurochemical process. Current approaches to sarcopenia focus primarily on skeletal muscle, whereas our findings support the concept that targeting neuromodulatory pathways upstream of the motor unit may represent an alternative strategy for preserving neuromuscular function. Although chemogenetic manipulation is not directly translatable, the noradrenergic system represents an attractive therapeutic target because its signaling pathways are pharmacologically accessible. In particular, our pharmacological experiments identify α1-adrenergic receptor signaling as a principal mediator of the excitatory effects of endogenous NE on motor neurons, suggesting that modulation of this pathway may provide a strategy to reduce maladaptive motor neuron excitation while preserving motor unit integrity during aging. Together, these observations raise the possibility that interventions aimed at restoring noradrenergic homeostasis could help preserve motor unit integrity and delay the progression of sarcopenia.

The present study has limitations. Although our findings strongly support a role for spinal noradrenergic overdrive in motor unit aging, future studies should determine whether chemogenetic suppression of descending noradrenergic activity normalizes motor neuron excitability in aged animals. Because the present study prioritized comprehensive histological evaluation of motor neurons, neuromuscular junctions and skeletal muscle, addressing this question will require independent experimental cohorts dedicated to electrophysiological analyses. Future studies should also determine the downstream cellular mechanisms by which α1-adrenergic receptor activation promotes motor unit deterioration, as well as the contributions of α2- and β-adrenergic signaling to spinal aging. In addition, it will be important to determine whether pharmacological restoration of noradrenergic homeostasis can preserve motor unit integrity and improve physical performance.

In summary, we identify spinal noradrenergic overdrive as a central mechanism linking aging to loss of motor unit integrity, motor neuron hyperexcitability, neuromuscular junction deterioration and skeletal muscle atrophy. By demonstrating that suppression of descending noradrenergic activity improves structural features of all major components of the aged motor unit, our findings shift the conceptual framework from one of inevitable neuromuscular degeneration to one in which important aspects of motor unit aging remain reversible. These results establish spinal noradrenergic signaling as a modifiable driver of motor unit aging and identify α1-adrenergic receptor-dependent motor neuron excitation as a potential therapeutic mechanism. Restoring noradrenergic homeostasis may therefore represent a strategy to preserve mobility, maintain motor unit integrity and attenuate sarcopenia in older individuals.

## Ethical guidelines

All experimental procedures were conducted in accordance with U.S. National Institutes of Health guidelines for the care and use of laboratory animals, and protocols A21-172 and A24-111 were reviewed and approved by the WFUSM Institutional Animal Care and Use Committee.

