## Supplementary Information for "Spinal Noradrenergic Overdrive Promotes Reversible Motor Unit Aging and Sarcopenia"

#### Supplementary Methods

##### Lumbar Spinal Cord Preparation for Motor Neuron Patch-Clamp Recording

Mice were deeply anesthetized with isoflurane and positioned in a stereotaxic frame equipped with three spinal adaptors (Stoelting, Wood Dale, IL, USA) on each side of the lumbar region. A laminectomy was performed at the T12–L1 vertebral levels to expose the L4–L5 spinal cord segments. Following decapitation, the spinal cord was rapidly extracted en bloc, and the meninges and associated blood were carefully removed. The tissue was then immediately immersed in ice-cold *dissecting and cutting* artificial cerebrospinal fluid (aCSF) containing (in mM): 0.75 K-gluconate, 1.25 $KH_2PO_4$, 20 glucose, 191 sucrose, 4 $MgCl_2 \cdot 7H_2O$, 1 $CaCl_2$, 26 choline bicarbonate, 1 Na-ascorbate, 5 ethyl pyruvate, and 3 myo-inositol. The temperature of the dissecting aCSF was maintained at approximately 4°C, while the cutting aCSF was kept at ~1°C using a Huber Minichiller 300 (Raleigh, NC, USA). The solution was adjusted to pH 7.4 with an osmolarity of 320 mOsm/kg.

The spinal cord was embedded in 5% (wt/vol) low-gelling-temperature agarose (Sigma Millipore, St. Louis, MO), allowed to gel at room temperature, and affixed to the stage of a vibratome (Leica VT1200S, Wetzlar, Germany). Oblique slices of 270-300 µm thickness were prepared and incubated at 35°C for 30 minutes in *recording* aCSF containing (in mM): 119 NaCl, 2.5 KCl, 120 $NaH_2PO_4$, 10 glucose, 25 $NaHCO_3$, 2 $CaCl_2$, 1 $MgCl_2$, 0.5 Na-ascorbate, 3

Na-pyruvate, and 3 myo-inositol, with an osmolarity of 315 mOsm/kg.[1] The solution was continuously bubbled with a gas mixture of 95% $O_2$ and 5% $CO_2$ to maintain a pH of 7.4. Slices were then allowed to recover at room temperature for approximately 30 minutes prior to electrophysiological recordings.

Recordings were performed in a custom-built chamber that replaced the stage of an Olympus BX51WI microscope (Tokyo, Japan). Slices were superfused with recording solution at a flow rate of 3–4 ml/min.

**Motor Neuron Whole-Cell Patch-Clamp Recording**

Whole-cell patch-clamp recordings of motor neurons were performed using established protocols previously applied to other neuronal types.[2] High-resistance seals (>2 GΩ) were achieved using infrared differential interference contrast (IR-DIC) optics (DAGE-MTI, IR-2000, Michigan City, IN). Patch electrodes were fabricated from borosilicate glass with filament (Sutter Instruments, Novato, CA, USA) using a DMZ Zeitz puller (Munich, Germany), yielding tip resistances of 3–6 MΩ. Electrodes were filled with an internal solution containing (in mM): 135 K-gluconate, 5 KCl, 2 $MgCl_2$, 10 HEPES, 0.2 EGTA, 2 Mg-ATP, 2 $Na_2$-GTP, and 1 Lucifer Yellow (Sigma Millipore). The solution was adjusted to pH 7.3 with KOH and had an osmolarity of 300 mOsm/L. Current-clamp recordings were conducted using a Multiclamp 700B amplifier and Clampex 10.7 software (Molecular Devices, San José, CA, USA). Electrode capacitance was compensated, and recordings were discarded if series resistance increased by more than 20% during the session. Signals were filtered at 1–4 kHz and digitized at 10–20 kHz using a Digidata 1440 interface (Molecular Devices). Data were analyzed using Clampfit 11.2 (Molecular Devices) and Excel 2016 (Microsoft, Redmond, WA, USA).

To confirm motor neuron identity, spinal cord slices were fixed overnight in 4% paraformaldehyde (PFA) at 4°C, rinsed in phosphate-buffered saline (PBS), and blocked overnight at 4°C in PBS containing 10% donkey serum and 1% Triton X-100. Slices were then incubated overnight at 4°C with an anti-choline acetyltransferase (ChAT) antibody (MilliporeSigma; Supplementary Table 1) diluted in PBS containing 1% donkey serum. Following washes in PBST, sections were incubated overnight at 4°C with Alexa Fluor 555-conjugated donkey anti-goat secondary antibody (Invitrogen, Thermo Fisher Scientific) diluted in PBS containing 1% donkey serum. The next day, sections were washed in PBST and counterstained with Hoechst 33342 (Thermo Fisher Scientific) for 5 min at room temperature, followed by a final PBS wash. Sections were mounted with fluorescence mounting medium (Dako) and imaged using an Olympus IX81 inverted fluorescence microscope equipped with an Orca-R2 CCD camera (Hamamatsu Photonics, Hamamatsu, Japan). Image acquisition was controlled using MetaMorph software (Molecular Devices).

**Skeletal Muscle Immunohistochemistry for Myofiber Cross-Sectional Area and Myosin Heavy-Chain Composition**

EDL and soleus muscles were embedded vertically in thickened optimal cutting temperature (O.C.T.) compound (Tissue-Tek, Sakura Finetek, Japan) mixed with baby powder in 35-mm dishes and rapidly frozen in liquid nitrogen. Muscles were cryosectioned transversely for the analysis of myosin heavy-chain (MyHC) composition. Myofiber classification was performed as previously described. [3],[4]

Cryosections were rinsed in PBS and blocked for 1 h in 10% goat serum in PBS. Myofiber subtypes were identified using antibodies against MyHC type I, type IIA and type IIB isoforms

(Supplementary Table 1). Sections were incubated with primary antibodies diluted in 10% goat serum in PBS for 2 h at room temperature. Following two 5-min washes in PBS, sections were incubated for 1 h at room temperature with the following secondary antibodies: goat anti-mouse IgG2b Alexa Fluor 350 (for BA-F8), goat anti-mouse IgG Alexa Fluor 488 (for SC-71), and goat anti-mouse IgM Alexa Fluor 555 (for BF-F3). All secondary antibodies were obtained from Invitrogen (Thermo Fisher Scientific, Carlsbad, CA, USA). This approach also identified hybrid fibers, which exhibited lower fluorescence intensity in multiple channels than pure fiber types.[4] Sections were mounted with fluorescence mounting medium (Dako S3023, Agilent Technologies, Carpinteria, CA, USA) and imaged using an Olympus IX81 inverted fluorescence microscope equipped with an Orca-R2 CCD camera (Hamamatsu Photonics). Image acquisition was controlled using MetaMorph software (Molecular Devices).

**Fast-Scan Cyclic Voltammetry In Vivo Recording**

Electrically evoked catecholamine release was recorded in the ventral horn of the lumbar spinal cord in anesthetized C57BL/6 mice. Mice were anesthetized with a single intraperitoneal injection of urethane (1.5 g/kg) and secured in a stereotaxic frame. Craniotomies and laminectomies were performed to permit electrode placement in the spinal cord and brainstem regions, specifically the pontine A5 group and the locus coeruleus (LC).

A bipolar stimulating electrode (Plastics One, Roanoke, VA, USA) was inserted into either the A5 group (coordinates: AP −5.3 mm, ML 1.3 mm, DV −5.1 to −5.2 mm) or the LC nucleus (coordinates: AP −5.3 mm, ML 0.75 mm, DV −3.2 mm).[5] A carbon fiber microelectrode was positioned in the ventral horn of the lumbar spinal cord at levels L3–L5 (ML 0.3–0.5 mm, DV

−1.0 to −1.4 mm), and an Ag/AgCl reference electrode was implanted contralaterally within the spinal cord.

Electrodes were connected to a voltammetric amplifier interfaced with a computer running Tar Heel CV 4.8 software (Chapel Hill, NC, USA). Catecholamine release was detected at the carbon fiber electrode every 100 ms using a triangular voltage waveform (−0.4 V to +1.3 V and back to −0.4 V vs. Ag/AgCl) applied at a scan rate of 400 V/s and a frequency of 10 Hz, as previously described[4]. Signals were validated by background-subtracted cyclic voltammograms, characterized by oxidation and reduction peaks at approximately +0.65 V and −0.2 V, respectively.

To assess frequency-dependent NE release in adult versus aged mice, electrical stimulation of the LC was delivered at varying frequencies (5, 10, 20, and 40 Hz) using 1-second pulses at 300 µA, with a 10-minute interval between stimulations.

**Fast-Scan Cyclic Voltammetry: In Vitro Recording**

Mice were anesthetized with a single i.p. injection of urethane prior to decapitation, and their spinal cords were rapidly excised and placed in ice-cold oxygenated artificial cerebrospinal fluid (aCSF) containing (in mM): 126 NaCl, 2.5 KCl, 1.2 $NaH_2PO4$, 2.4 $CaCl_2$, 1.2 $MgCl_2$, 0.4 L-ascorbic acid, 11 glucose, and 25 $NaHCO_3$, adjusted to pH 7.40. Coronal slices (350 µm thick) of the L4-L5 spinal cord segments were prepared using a vibrating tissue slicer (Vibratome 1000 Plus, The Vibratome Company, St. Louis, MO, USA), and allowed to equilibrate for 30 minutes in oxygenated aCSF at room temperature.

Slices were transferred to a submersion recording chamber perfused with oxygenated aCSF flowing at a flow rate of 1 mL/min. A twisted bipolar stimulating electrode (Plastics One, Roanoke, VA, USA) was positioned in the ventral spinal cord targeting Rexed laminae VIII-IX. A carbon fiber recording electrode, connected to a voltammetric amplifier (UNC Electronics Design Facility, Chapel Hill, NC, USA), was placed adjacent to both terminals of the stimulation electrode, approximately 300 μm apart.

NE release was evoked by a 1-second electrical stimulation (330 μA, 2 ms/phase, monophasic) at various frequencies (5, 10, 20, 30, and 40 Hz) every 10 minutes. Extracellular NE was recorded every 100 ms for 15 seconds using a triangular waveform (-0.4 V to +1.3 V and back to -0.4 V vs Ag/AgCl) at a scan rate of 400V/s. NE was identified by background-subtracted cyclic voltammograms showing characteristic oxidation and reduction peaks.

***Microdialysis Measurements of Extracellular NE Concentration***

Extracellular NE concentration was measured using ventral lumbar spinal cord microdialysis. Mice were anesthetized with a single i.p. injection of urethane and secured in a stereotaxic frame. A hole was drilled to facilitate the insertion of a microdialysis probe (1 mm membrane length, 0.24 mm diameter, Cuprophane, 6 kDa cut-off, CMA-7, Harvard Apparatus, Holliston, MA, USA). The stereotaxic coordinates for the implantation were the same as those used for the insertion of the working electrode in voltammetric recordings. Following implantation, the microdialysis probe was connected to a CMA 402 syringe pump (Harvard Apparatus) and perfused with artificial cerebrospinal fluid (aCSF) containing (in mM): 150 $Na^+$, 3.0 $K^+$, 1.4 $Ca^{2+}$, 0.8 $Mg^{2+}$, 1.0 $PO_4^{3-}$, and 155 $Cl^-$ (Harvard Apparatus, Holliston, MA, USA). After an equilibration period consisting of perfusion at 1 μl $min^{-1}$ for 1 h followed by 0.2 μl $min^{-1}$ for an additional 1 h, 20 μl dialysate samples were collected. A volume of 5 μl of 4% acetic acid was added immediately to each sample. Dialysates were subsequently analyzed for NE content using high-performance liquid chromatography with electrochemical detection HPLC-ECD (Amuza, San Diego, CA, USA).

**Ventral Spinal Cord NE Content**

For quantification of NE content, the L4–L5 spinal cord segments were embedded in agar as previously described, and the ventral portion was carefully separated from the dorsal cord. Ventral hemisections were immediately flash-frozen in liquid nitrogen and stored at −80°C in cryovials until further processing.

Tissue samples were manually homogenized in 0.1 M perchloric acid ($HClO_4$), followed by centrifugation at 12,000 rpm for 5 minutes at 4°C. The resulting supernatant was filtered through 0.22 µm syringe filters and used for NE quantification via HPLC-ECD; Amuza). Protein concentration was determined using the bicinchoninic acid (BCA) assay (Pierce BCA Protein Assay Kit, Thermo Fisher Scientific, Rockford, IL, USA).

Monoamines were separated using a reverse-phase column (EICOMPAK CA-SODS, 2.1 mm ID × 150 mm) with a mobile phase composed of 95% 0.1 M sodium phosphate buffer (pH 6.0) containing 400 mg/L sodium octanesulfonate, 50 mg/L EDTA, and 5% methanol. The flow rate was maintained at 230 µL/min. NE was detected using a graphite working electrode (WE-3G, EiCOM) set at +450 mV.

**Western Blot and Protein Expression Analysis**

Western blotting was performed to quantify the expression of the NE transporter (NET) and α2A-adrenergic receptor in ventral lumbar spinal cord tissue. Tissue samples were homogenized in Laemmli sample buffer, heated at 100°C for 4 min and stored at −20°C until analysis.

Protein lysates were separated on Criterion TGX 8–16% precast gels (Bio-Rad, Hercules, CA, USA) using SDS–PAGE and transferred onto PVDF membranes. Membranes were blocked for 1 h at room temperature in 5% non-fat dry milk prepared in TBST (20 mM Tris-HCl, pH 7.5, 500 mM NaCl and 0.1% Tween-20).

Membranes were incubated overnight at 4°C with rabbit anti-α2A-adrenergic receptor (Thermo Fisher Scientific), rabbit anti-NET (Sigma-Aldrich) and mouse anti-β-actin (Santa Cruz Biotechnology) antibodies diluted in blocking buffer (Supplementary Table 1). After three washes in TBST, membranes were incubated for 1 h at room temperature with horseradish peroxidase-conjugated goat anti-rabbit IgG or goat anti-mouse IgG secondary antibodies. Immunoreactive bands were visualized using enhanced chemiluminescence and imaged with a Bio-Rad ChemiDoc imaging system.

For protein normalization, membranes were stripped using Restore PLUS Western Blot Stripping Buffer (Thermo Fisher Scientific), re-blocked and reprobed with anti-β-actin antibody. Densitometric analysis was performed using Image Lab v6.1 software (Bio-Rad). Lane alignment and background subtraction were applied uniformly across all samples. Adjusted band volumes were exported for quantitative analysis.

**Supplementary Figures**

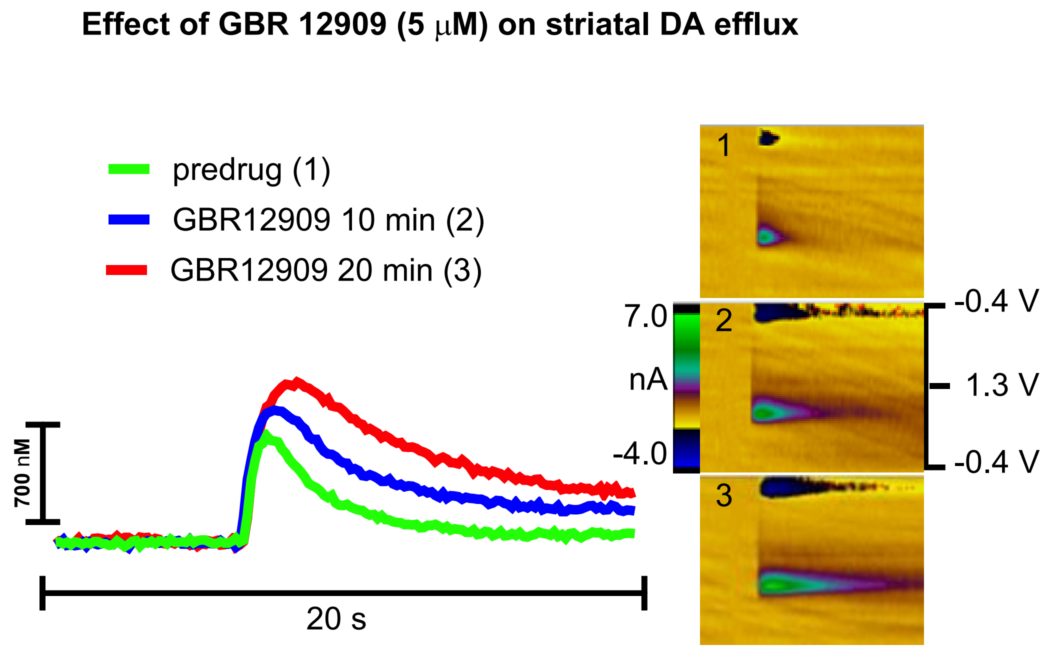

**Supplementary Figure 1**

**Supplementary Figure 1 (GBR).** ***GBR 12909 enhances electrically evoked dopamine release in striatal slices.*** Representative traces (left) and corresponding 2D pseudocolor plots (right) illustrate electrochemical changes in the dorsal striatum of mouse brain slices before (1), and after 10 (2) and 20 (3) minutes of exposure to GBR 12909 (5 μM). The pseudocolor plots display voltammetric data with time on the x-axis, applied scan potential on the y-axis, and background-subtracted faradaic current on the z-axis. The green signal indicates oxidation currents corresponding to dopamine release.

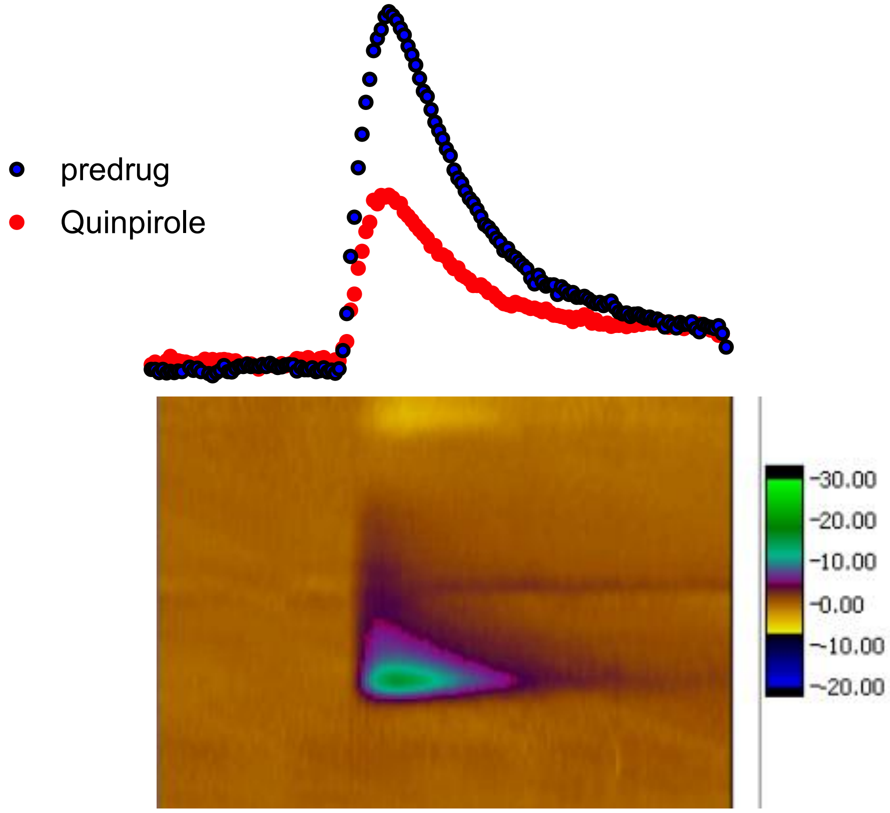

**Supplementary Figure 2**

**Supplementary Figure 2.** ***Quinpirole reduces electrically evoked dopamine release in striatal slices.*** Representative traces (top) and corresponding 2D pseudocolor plots (bottom) illustrate electrochemical changes recorded in real time in the dorsal striatum of mouse brain slices before (red) and after 20 minutes of quinpirole (10 µM, black) perfusion. The pseudocolor plots display voltammetric data with time on the x-axis, applied scan potential on the y-axis, and background-subtracted faradaic current on the z-axis. The green signal indicates oxidation currents corresponding to dopamine release.

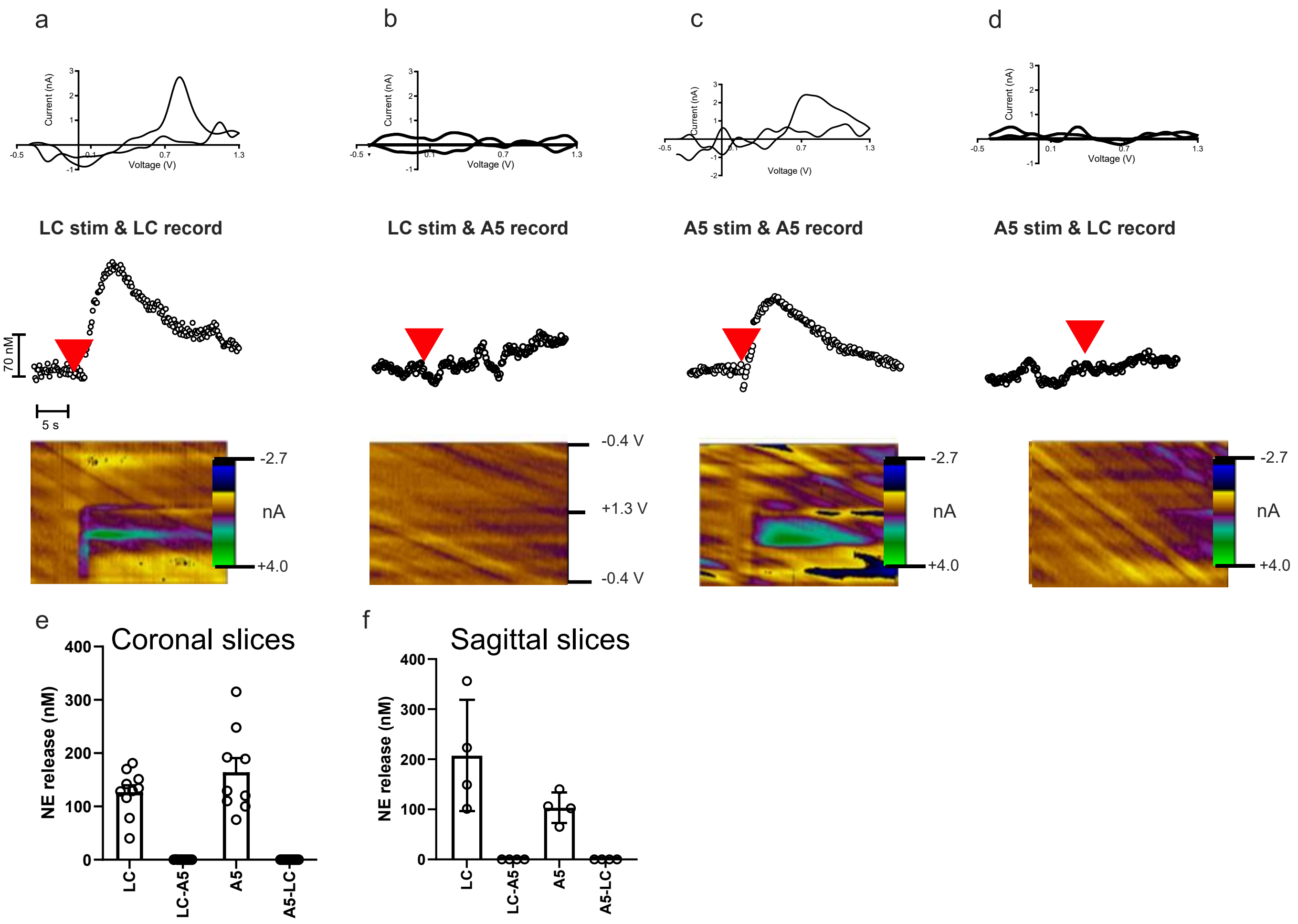

**Supplementary Figure 3**

**Supplementary Figure 3.** ***Functional connectivity between the ventrolateral pons encompassing the A5 noradrenergic cell group and the locus coeruleus in coronal and sagittal brainstem slices.*** Fast-scan cyclic voltammetry (FSCV) recordings of norepinephrine (NE) release evoked by electrical stimulation of the locus coeruleus (LC) or the ventrolateral pontine A5 noradrenergic region. a, LC stimulation with FSCV recording in the LC. b, LC stimulation with FSCV recording in the A5 region. c, A5 region stimulation with FSCV recording in the A5 region. d, A5 region stimulation with FSCV recording in the LC. Top panels show representative cyclic voltammograms identifying NE; middle panels show extracellular NE concentration as a function of time, with red arrowheads indicating stimulation onset; bottom panels show pseudocolor plots of current as a function of applied potential and time. Peak NE release was quantified for each stimulation-recording configuration. e**,** Peak NE release quantified for each stimulation-recording configuration. Mann-Whitney analyses demonstrated significant local NE release following LC stimulation recorded in the LC (LC→LC vs LC→A5, $P = 0.0004$) and following A5 stimulation recorded in the A5 region (A5→A5 vs A5→LC, $P = 0.0002$). No detectable NE release was observed in cross-nucleus recording configurations, indicating functional independence of LC and A5 noradrenergic signaling pathways. f, Mann-Whitney analyses demonstrated significant local NE release following LC stimulation recorded in the LC (LC→LC vs LC→A5, $P = 0.0286$) and following A5 stimulation recorded in the A5 region (A5→A5 vs A5→LC, $P = 0.0286$). No detectable NE release was observed in cross-nucleus recording configurations, indicating functional independence of LC and A5 noradrenergic signaling pathways. Data are presented as mean ± s.e.m., with individual data points representing independent recordings.

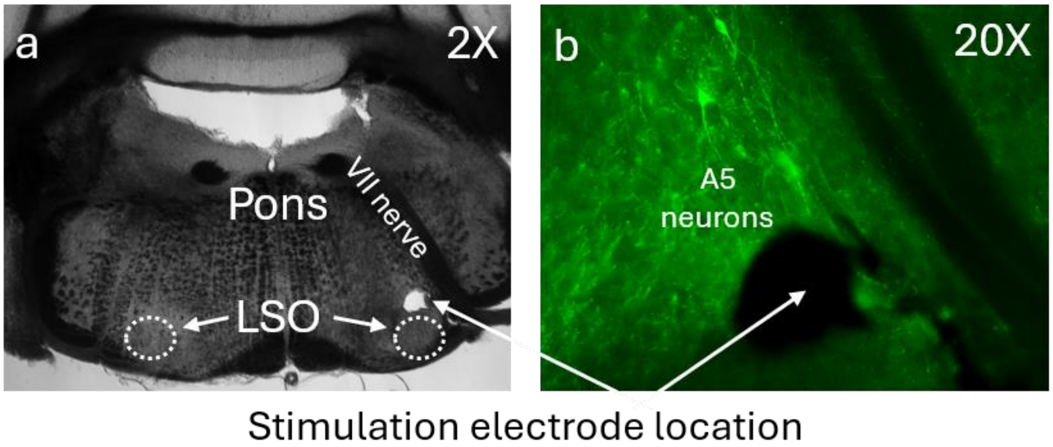

**Supplementary Figure 4**

**Supplementary Figure 4. Electrode placement for stimulation of the ventrolateral pontine region containing the A5 noradrenergic cell group.** a, Low-magnification brightfield image (2×) of a 100-µm-thick coronal pontine section from a young (4-month-old) mouse showing the fourth ventricle, facial nerve (cranial nerve VII), and lateral superior olive (LSO; dotted outlines). b, Higher-magnification image (20×) of the same section immunolabeled for tyrosine hydroxylase, illustrating the location of the stimulation electrode relative to A5 noradrenergic neurons within the ventrolateral pontine region.

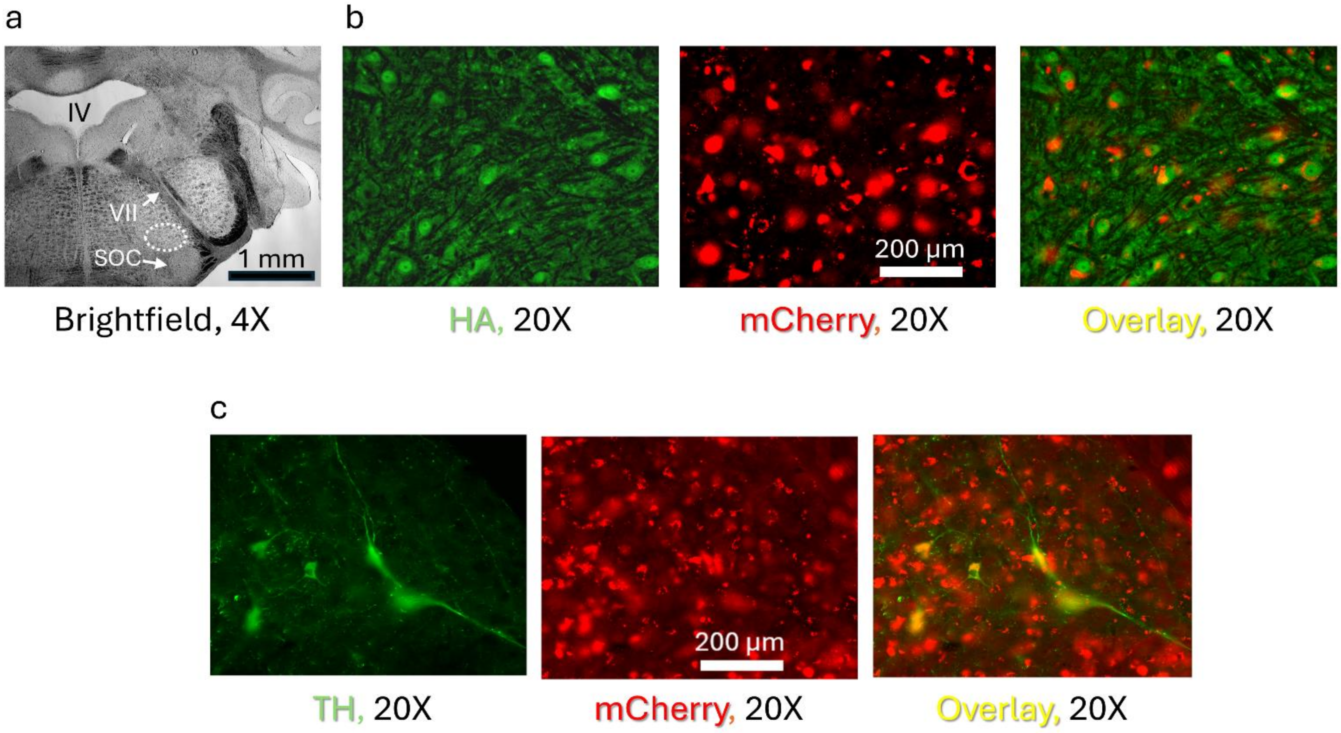

**Supplementary Figure 5**

**Supplementary Figure 5. Retrograde expression of inhibitory DREADDs in spinally projecting noradrenergic neurons of the pontine A5 region.** (a) Low-magnification brightfield image of a coronal brainstem section showing the anatomical location of viral expression within the ventrolateral pontine region containing the A5 noradrenergic cell group. The outlined area indicates the region analyzed for viral transduction. IV, fourth ventricle; VII, facial motor nucleus; SOC, superior olivary complex. Scale bar = 1 mm. (b) Representative fluorescence images of the A5 region demonstrating expression of the retrogradely transported CAV2-PRS-HA-hM4Di(E1)-hSyn-mCherry(3) vector. Hemagglutinin (HA) immunoreactivity (green), mCherry fluorescence (red), and the merged image confirm expression of the hM4Di-mCherry construct within the targeted pontine neurons. Scale bar = 200 µm. (c) Representative images showing tyrosine hydroxylase (TH) immunolabeling (green), mCherry fluorescence (red), and the merged image in the ventrolateral pontine A5 region. Colocalization of TH and mCherry demonstrates transduction of catecholaminergic neurons that project to the lumbar spinal cord. Scale bar = 200 µm. These data confirm successful retrograde targeting of spinally projecting noradrenergic neurons in the A5 pontine noradrenergic nucleus following lumbar spinal cord injection of CAV2-PRS-HA-hM4Di(E1)-hSyn-mCherry(3).

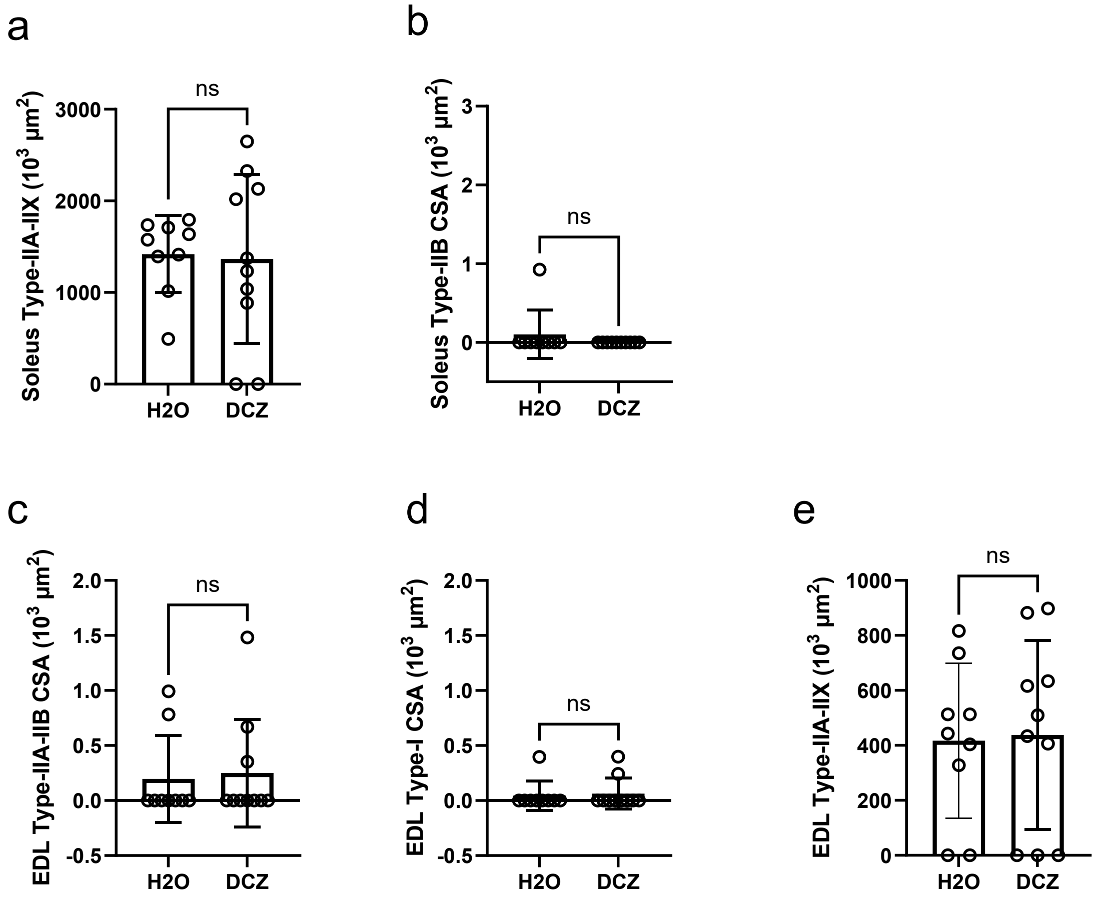

**Supplementary Figure 6**

**Supplementary Figure 6. Cross-sectional area of infrequent muscle fiber subtypes in aged mice following chemogenetic modulation of the A5 noradrenergic system.** a,b, Cross-sectional area (CSA) of soleus type-IIA/IIX (a) and type-IIB (b) fibers in mice receiving drinking water ($H_2O$) or deschloroclozapine (DCZ). c–e, CSA of extensor digitorum longus (EDL) type-IIA/IIB (c), type-I (d), and type-IIA/IIX (e) fibers. No significant differences were detected between treatment groups for any fiber subtype analyzed. Bars represent mean ± s.d., and circles represent individual mice. Statistical comparisons were performed using two-tailed unpaired *t*-tests. ns, not significant.

#### Supplementary Table 1

#### Antibodies and Toxin used for Immunohistochemistry and Western Blot

| Antibody | Dilution | Source (Catalog No.) |
|---|---|---|
| **Skeletal Muscle** | | |
| Laminin | 1:100 | Millipore Sigma (MAB1914P) |
| Goat anti-rat IgG Alexa Fluor-680 | 1:500 | Invitrogen, Thermo Fisher (A21096) |
| MyHC-Type I | 1:50 | DBHS (Developmental Studies Hybridoma Bank-Univ Iowa). (BA-F8-c) |
| Goat anti-mouse IgG 2b Alexa Fluor-350 | 1:500 | Invitrogen, Thermo Fisher (A21140) |
| MyHC-Type IIa | 1:500 | DBHS (SC-71-c) |
| Goat anti-mouse IgG Alexa Fluor-488 | 1:500 | Invitrogen, Thermo Fisher (A21121) |
| MyHC-Type IIb | 1:100 | DBHS (BF-F3-c) |
| Goat anti-mouse IgM Alexa Fluor-555 | 1:500 | Invitrogen, Thermo Fisher (A21426) |
| **Neuromuscular Junction** | | |
| SV2 | 1:25 | DBHS (SV2-c) |
| Neurofilament SMI 311 | 1:500 | Biolegend (837801) |
| Goat anti-mouse IgG Alexa Fluor-488 | 1:500 | Invitrogen, Thermo Fisher (A21121) |
| α-Bungarotoxin - tetramethylrhodamine conjugated | 1:250 | Biotium (T1175) |
| **Spinal Cord** | | |
| ChAT | 1:100 | Millipore Sigma (AB144P) |
| Donkey anti-goat Alexa Fluor 488 | 1:500 | Invitrogen, Thermo Fisher (A11055) |
| **Western Blot** | | |
| α-A2 AR | 1:600 | ThermoFisher (PA1-048) |
| Norepinephrine Transporter (NET) | 1:300 | Millipore Sigma (AB5066P) |
| β-actin | 1:600 | Santa Cruz (S47778) |
| **Brainstem** | | |
| Tyrosine hydroxylase | 1:100 | Millipore Sigma (AB-152) |
| Hemagglutinin | 1:100 | Novus Biologicals ( NB600-363) |
| Donkey anti-rabbit IgG Alexa Fluor-488 | 1:500 | Invitrogen, Thermo Fisher (A21206) |
| Hoechst | 1:2000 | Thermo Fisher |

#### Supplementary Table 2. Pharmacological agents used in the study

| Agent | Manufacturer (Catalog No.) | Manufacturer location | Concentration |
|---|---|---|---|
| BRL 44408 maleate | Tocris (1133) | Avonmouth, Bristol, UK | 2 µM |
| Clonidine hydrochloride | Sigma-Aldrich (C7897) | St. Louis, MO, USA | 10 µM |
| Cocaine | The National Institute on Drug Abuse (NIDA) | Bethesda, MD, USA | 10 µM |
| Deschloroclozapine | Tocris (7193) | Avonmouth, Bristol, UK | 10 µg $ml^{-1}$ |
| Desipramine hydrochloride | Tocris (7193) | Avonmouth, Bristol, UK | 10 µM |
| Fluoxetine hydrochloride | Millipore-Sigma (F132) | Burlington, MA, USA | 10 µM |
| GBR12909 dehydrochloride | Tocris (0421) | Avonmouth, Bristol, UK | 5 µM |
| Idazoxan hydrochloride | Sigma MKCG 6225 | St. Louis, MO, USA | 10 µM |
| Isoflurane | Piramal Pharma Solutions (66794-013) | Lexington, KY, USA | 1.5-3% |
| Prazosin hydrochloride | Tocris (0623) | Avonmouth, Bristol, UK | 10 µM |
| Propranolol hydrochloride | Tocris (0624) | Avonmouth, Bristol, UK | 10 µM |
| Quinpirole hydrochloride | Tocris (1061) | Avonmouth, Bristol, UK | 10 µM |
| Raclopride | Tocris (1810) | Avonmouth, Bristol, UK | 10 µM |
| Reserpine | Tocris (2742) | Avonmouth, Bristol, UK | 10 µM |
| Urethane | Millipore-Sigma (U2500) | Burlington, MA, USA | 1.5 g/kg |
| Yohimbine hydrochloride | Tocris (1127) | Avonmouth, Bristol, UK | 10 µM |

### Supplementary Table 3. Statistical analyses, sample sizes, and *P* values

| Figure | Statistical test | n samples/n mice | *P* value |
|---|---|---|---|
| Fig. 1b. Membrane Potential | Linear regression | 135 MNs / 81 mice | <0.0001 |
| Fig. 1b. Membrane Capacitance | Linear regression | 135 MNs / 81 mice | <0.0001 |
| Fig. 1b. Input Resistance | Linear regression | 135 MNs / 81 mice | <0.0001 |
| Fig. 1b.Cumulative Frequency | Linear regression | 135 MNs / 81 mice | <0.0001 |
| Fig. 1b.AP Threshold | Linear regression | 135 MNs / 81 mice | 0.0036 |
| Fig. 2c. AP delay | Paired t-test, two tailed | 7 MNs / 7 mice | 0.0001 |
| Fig. 2c. Current elicit AP | Paired t-test, two tailed | 7 MNs / 7 mice | 0.0028 |
| Fig. 2d. Post-Opto/Pre-Opto Frequency Change | One-way ANOVA followed by Dunnett's multiple-comparisons test. | Untreated: 6 MNs / 6 mice;<br>Prazosin: 6 MNs / 6 mice;<br>Yohimbine: 6 MNs / 6 mice<br>BRL 44408: 6 MNs / 6 mice<br>Propranolol: 6 MNs / 6 mice | Prazosin: 0.0313<br>Yohimbime: >0.9999<br>BRL 44408: 0.996<br>Propranolol: 0.866 |
| Fig. 3c. Cocaine | Paired t-test, two tailed | 6 mice | 0.0154 |
| Fig. 3c. Reserpine | Paired t-test, two tailed | 6 mice | 0.0016 |
| Fig. 3c. GBR12909 | Paired t-test, two tailed | 8 mice | 0.2108 |
| Fig. 3c. Quinpirole | Unpaired t-test with Welch's correction | 6 mice | 0.8780 |
| Fig. 3c. Raclopride | Paired t-test, two tailed | 9 mice | 0.0954 |
| Fig. 3c. Fluoxetine | Paired t-test, two tailed | 8 mice | 0.2524 |
| Fig. 3c. Desipramine | Paired t-test, two tailed | 9 mice | 0.5101 |
| Fig. 3c. Idazoxan | Paired t-test, two tailed | 5 mice | 0.0299 |
| Fig. 3c. Clonidine | Paired t-test, two tailed | 8 mice | 0.0258 |
| Fig. 4a | Two-way ANOVA Sidak multiple comparisons | 8 young mice / 8 old mice for all 5 frequencies | 5Hz:0.9795<br>10Hz:0.6087<br>20Hz:0.0223<br>30Hz:0.0022<br>40Hz:<0.0001 |
| Fig. 4b | Unpaired t-test with Welch's correction | 16 young slices / 9 old slices | 0.0349 |
| Fig. 4d | Unpaired t-test with Welch's correction | 9 young slices / 7 old slices | 0.0016 |
| Fig. 5a. Extracellular NE | Unpaired t-test, two-tailed | 5 mice | 0.3257 |
| Fig. 5a. Extracellular MHPG | Unpaired t-test, two-tailed | 5 mice | 0.5997 |
| Fig. 5a. NE tissue content | Unpaired t-test, two-tailed | 5 mice | 0.0403 |
| Fig. 5a. MHPG tissue content | Unpaired t-test, two-tailed | 5 mice | 0.2589 |
| Fig. 5a. NM tissue content | Unpaired t-test, two-tailed | 5 mice | 0.5571 |
| Fig. 5a. DA tissue content | Unpaired t-test, two-tailed | 4 mice | 0.4020 |
| Fig. 5c. | Unpaired t-test, two-tailed | 10 mice | 0.0009 |
| Fig. 5e. | Unpaired t-test, two-tailed | 10 mice | 0.0046 |
| Fig. 6d. Mouse weight | Unpaired t-test with Welch's correction | H2O group: 9; DCZ group: 10 | 0.2991 |
| Fig. 6d. EDL/BW | Unpaired t-test with Welch's correction | H2O group: 9; DCZ group: 10 | 0.0111 |
| Fig. 6d. Soleus/BW | Unpaired t-test with Welch's correction | H2O group: 9; DCZ group: 10 | 0.3029 |
| Fig. 6d. Tibial Anterior/BW | Unpaired t-test with Welch's correction | H2O group: 9; DCZ group: 10 | 0.0344 |
| Fig. 6d. Gastrocnemius/BW | Unpaired t-test with Welch's correction | H2O group: 9; DCZ group: 10 | 0.0228 |
| Fig. 6d. EDL Total CSA | Unpaired t-test with Welch's correction | H2O group: 9; DCZ group: 10 | 0.0085 |
| Fig. 6d. EDL Laminin EC Space | Unpaired t-test with Welch's correction | H2O group: 9; DCZ group: 10 | 0.9965 |
| Fig. 6d. Soleus Total CSA | Unpaired t-test with Welch's correction | H2O group: 9; DCZ group: 10 | 0.7597 |
| Fig. 6d. Soleus Laminin EC Space | Unpaired t-test with Welch's correction | H2O group: 9; DCZ group: 10 | 0.1009 |
| Fig. 6d. Soleus Type-I CSA | Unpaired t-test with Welch's correction | H2O group: 9; DCZ group: 10 | >0.999 |
| Fig. 6d. Soleus Type-II CSA | Unpaired t-test with Welch's correction | H2O group: 9; DCZ group: 10 | 0.0335 |
| Fig. 6d. EDL Type-IIA CSA | Unpaired t-test with Welch's correction | H2O group: 9; DCZ group: 10 | 0.0165 |
| Fig. 6d. EDL Type-IIB CSA | Unpaired t-test with Welch's correction | H2O group: 9; DCZ group: 10 | 0.0078 |
| Fig. 6d. EDL Type-IIX CSA | Unpaired t-test with Welch's correction | H2O group: 9; DCZ group: 10 | 0.0467 |
| Fig. 6d. EDL IIX-IIB CSA | Unpaired t-test with Welch's correction | H2O group: 9; DCZ group: 10 | 0.0196 |
| Fig. 6d. EDL Type-IIB (%) | Unpaired t-test with Welch's correction | H2O group: 9; DCZ group: 10 | 0.7199 |
| Fig. 6d. EDL Type-IIB (#) | Unpaired t-test with Welch's correction | H2O group: 9; DCZ group: 10 | 0.1698 |
| Fig. 6d. EDL Type-IIB population (%) | Unpaired t-test with Welch's correction | H2O group: 9; DCZ group: 10 | 0.7038 |
| Fig. 7b. EDL NMJ Volume | Unpaired t-test with Welch's correction | H2O group: 9; DCZ group: 10 | 0.2545 |
| Fig. 7b. EDL NMJ Fragments | Unpaired t-test with Welch's correction | H2O group: 9; DCZ group: 10 | 0.5142 |
| Fig. 7b. EDL NMJ Manders tM1 | Unpaired t-test with Welch's correction | H2O group: 9; DCZ group: 10 | 0.0495 |
| Fig. 7b. EDL NMJ Manders tM2 | Unpaired t-test with Welch's correction | H2O group: 9; DCZ group: 10 | 0.8205 |
| Fig. 7b. EDL Denervation Index | Mixed-effects model followed by Sidak's multiple-comparisons test | H2O: n = 9; DCZ: n = 10 | FI:0.343<br>EPD:0.107<br>APD:0.214<br>FD:0.021 |

| | | | |
|---|---|---|---|
| Fig. 7d. SOL NMJ Volume | Unpaired t-test with Welch's correction | H2O group: 9; DCZ group: 10 | 0.0867 |
| Fig. 7d. SOL NMJ Fragments | Unpaired t-test with Welch's correction | H2O group: 9; DCZ group: 10 | 0.9103 |
| Fig. 7d. SOL NMJ Manders tM1 | Unpaired t-test with Welch's correction | H2O group: 9; DCZ group: 10 | 0.5043 |
| Fig. 7d. SOL NMJ Manders tM2 | Unpaired t-test with Welch's correction | H2O group: 9; DCZ group: 10 | 0.9832 |
| Fig. 7d. SOL NMJ Denervation Index | Mixed-effects model followed by Sidak's multiple-comparisons test | H2O: n = 9; DCZ: n = 10 | FI:0.530<br>EPD:0.240<br>APD:0.450<br>FD:0.290 |
| Fig. 7e. MN Soma Area | Unpaired t-test with Welch's correction | H2O group: 9; DCZ group: 10 | 0.0353 |
| Fig. 7e. Number of MN / VH | Unpaired t-test with Welch's correction | H2O group: 9; DCZ group: 10 | 0.0591 |
| Supplementary Fig. 3e | Mann-Whitney analysis | LC: 10; A5 zone: 9; LC-A5: 10<br>A5-LC: 9 | LC→LC vs LC→A5, *P* = 0.0004<br>A5→A5 vs A5→LC, *P* = 0.0002 |
| Supplementary Fig. 3f | Mann-Whitney analysis | LC: 4; A5 zone: 4; LC-A5: 4<br>A5-LC: 4 | LC→LC vs LC→A5, *P* = 0.0286<br>A5→A5 vs A5→LC, *P* = 0.0286 |
| Supplementary Fig. 6a | Unpaired t-test with Welch's correction | H2O group: 9; DCZ group: 10 | 0.8731 |
| Supplementary Fig. 6b | Unpaired t-test with Welch's correction | H2O group: 9; DCZ group: 10 | 0.3466 |
| Supplementary Fig. 6c | Unpaired t-test with Welch's correction | H2O group: 9; DCZ group: 10 | 0.7955 |
| Supplementary Fig. 6d | Unpaired t-test with Welch's correction | H2O group: 9; DCZ group: 10 | 0.7504 |
| Supplementary Fig. 6e | Unpaired t-test with Welch's correction | H2O group: 9; DCZ group: 10 | 0.8863 |
